# Transcriptomic profiling identifies cellular and molecular transitions during Merkel cell regeneration

**DOI:** 10.64898/2026.09.16.752174

**Authors:** Ahlan S. Ferdous, Elgene J. A. Quitevis, Eric Peterman, Sheridan M. Sargent, Erik C. Black, Sarthak M. Dighe, Jeffrey P. Rasmussen

## Abstract

Skin exhibits a remarkable capacity for regeneration, which requires the coordinated reconstitution of diverse cell types and structures. Merkel cells are mechanosensory cells that detect gentle touch and develop in coordination with dermal appendages, yet Merkel cell regeneration remains incompletely understood. To investigate Merkel cell lineage dynamics during skin regeneration, we performed single-cell RNA sequencing of regenerating adult zebrafish skin after dermal appendage removal. We find that the regenerating Merkel cell lineage is heterogeneous and characterize molecular markers associated with Merkel cell subpopulations. Through cross-species analysis with mouse and human Merkel cells, we identify conserved elements of Merkel cell identity and function across vertebrates. Trajectory analysis reveals a continuous spectrum of transcriptional state transitions within the Merkel cell population, suggesting dynamic lineage progression during regeneration and involving an epithelial-to-mesenchymal-like transition. Finally, cell fate mapping and *in vivo* analysis identify a putative pool of *cldna+* suprabasal keratinocyte progenitors that give rise to regenerating Merkel cells. Together, these findings establish a genome resource for studies of zebrafish skin regeneration and provide insights into Merkel cell heterogeneity, conservation, lineage relationships and regenerative dynamics.

## INTRODUCTION

Skin protects our internal organs from physical injury, dessication and environmental insults. In addition to this barrier role, the skin’s sensory functions are essential for surviving our evolving physical environment, forming social bonds and promoting cognitive development (Jenkins and Lumpkin, 2017). Vertebrate skin contains epidermal and dermal layers composed of diverse cell types (e.g., epithelial, immune and sensory) and appendages (e.g., scales, feathers and hairs) that contribute to skin function and homeostasis (Chuong et al., 2025). Importantly, the skin has a high capacity for regeneration, allowing it to continuously renew itself and repair damage through coordinated cell proliferation and differentiation (Blanpain and Fuchs, 2009).

Specialized neuronal and sensory cells endow skin with remarkable sensitivity to environmental stimuli (Handler and Ginty, 2021). For example, mechanosensory epidermal cells known as Merkel cells (MCs) and associated somatosensory neuron endings form the bipartite MC-neurite complex (Hartschuh et al., 1986). The MC-neurite complex detects gentle touch inputs, including textures and curvatures (Maricich et al., 2009; Maricich et al., 2012). MC-neurite complex function requires that MCs express the mechanosensitive ion channel Piezo2 (Ikeda et al., 2014; Maksimovic et al., 2014; Woo et al., 2014) and release synaptic vesicles to modulate neuronal activity (Hoffman et al., 2018; Yamada et al., 2024). Genetic loss of MCs in mice can lead to abnormal responses to light touch, such as triggering itch-evoked behaviors (Feng et al., 2018) or pain perception (Jeon et al., 2021). MCs are further associated with Merkel cell carcinoma (MCC), an aggressive skin cancer (Harms et al., 2018). Thus, studies of MC development and regeneration inform our understanding of skin function and homeostasis.

Although MCs populate diverse types of vertebrate skin and oral mucosa (Hartschuh et al., 1986), the mechanisms of MC development and regeneration have primarily been studied in mouse hairy skin (Zhou and Ezhkova, 2025). In this system, MCs form touch domes, crescent-shaped groupings of cells around hair follicles, and develop from epidermal basal keratinocyte stem cell precursors (Morrison et al., 2009; Van Keymeulen et al., 2009). Genetic experiments in hairy skin implicate the neurogenic transcription factors Atoh1 (Atonal bHLH transcription factor 1) and Sox2 (SRY-box transcription factor 2) in MC development (Bardot et al., 2013; Perdigoto et al., 2014).

Mild injury to hairs triggers MC regeneration (Wright et al., 2017), yet the underlying mechanisms have not been thoroughly investigated. Challenges to studying MC regeneration include the opacity of mammalian skin and relative scarcity of MCs (Lacour et al., 1991).

Zebrafish MCs are optically accessible and share conserved cellular features with mammalian MCs such as a basal keratinocyte origin, somatosensory innervation and actin-rich microvilli (Brown et al., 2023). At the molecular level, zebrafish MCs express *piezo2*, neurosecretory molecules and orthologs of Atoh1 and Sox2 (Brown et al., 2023). Superficial damage to the trunk skin by dermal appendage removal triggers the *de novo* regeneration of zebrafish MCs (Craig et al., 2025). We previously used dermal appendage development and regeneration to identify the direct precursors of MCs, a transient cell population called dendritic Merkel cells (dMCs) due to their morphology (Craig et al., 2025). dMCs have been described in diverse types of human and rodent skin and oral mucosa (Kim and Holbrook, 1995; Moll et al., 1984; Nakafusa et al., 2006; Narisawa et al., 1993; Tachibana et al., 1997; Tachibana et al., 1998) and live-cell imaging in zebrafish revealed that dMCs exhibit dynamic migratory behaviors (Craig et al., 2025). Despite the identification of dMCs as MC precursors, the molecular and cellular trajectory of the MC lineage during regeneration remains unclear.

To capture the transcriptomic changes that accompany MC lineage regeneration, we performed single-cell RNA sequencing (scRNA-seq) on zebrafish trunk skin during dermal appendage regeneration. Our dataset encompasses regenerating dermal and epidermal cells, including MCs. We identified known and novel MC lineage markers that encompass diverse biological functions and phases of MC regeneration. We compared our dataset with published human and mouse MC transcriptomics to identify a core gene expression profile of vertebrate MCs. We identified a cluster of dMCs within the MC lineage and found them enriched in an epithelial-to-mesenchymal transition (EMT) signature, which we confirmed *in situ*. Finally, differentiation trajectory analysis and generation of a novel transgenic line indicated that *cldna+* suprabasal keratinocytes can give rise to the regenerating MC lineage. Our study thus provides a scRNA-seq transcriptomic resource for studies of skin regeneration and a molecular and cellular analysis of MC lineage progression.

## RESULTS

### Single-cell RNA-seq captures diverse regenerating dermal and epidermal cells

The adult zebrafish trunk skin contains a stratified epidermis that overlays scales, a type of dermal appendage formed by osteoblasts (**Fig. 1A**) (Aman and Parichy, 2024; Le Guellec et al., 2004). The epidermis comprises both keratinocyte and non-keratinocyte cell types (**Fig. 1A**). The keratinocyte compartment contains three molecularly and morphologically distinct layers: basal, suprabasal and superficial (Hou et al., 2020; Marí-Beffa et al., 1996). The non-keratinocyte compartment contains heterogeneous cell types, including mucus-secreting goblet cells, immune cells and MCs (Brown et al., 2023; Le Guellec et al., 2004; Lin et al., 2019; Peterman et al., 2023). Scales and associated epidermis rapidly regenerate following scale removal (Aman et al., 2018; Cox et al., 2018; Iwasaki et al., 2018; Rasmussen et al., 2018; Santoso et al., 2024; Sire et al., 1997).

**Figure 1.**
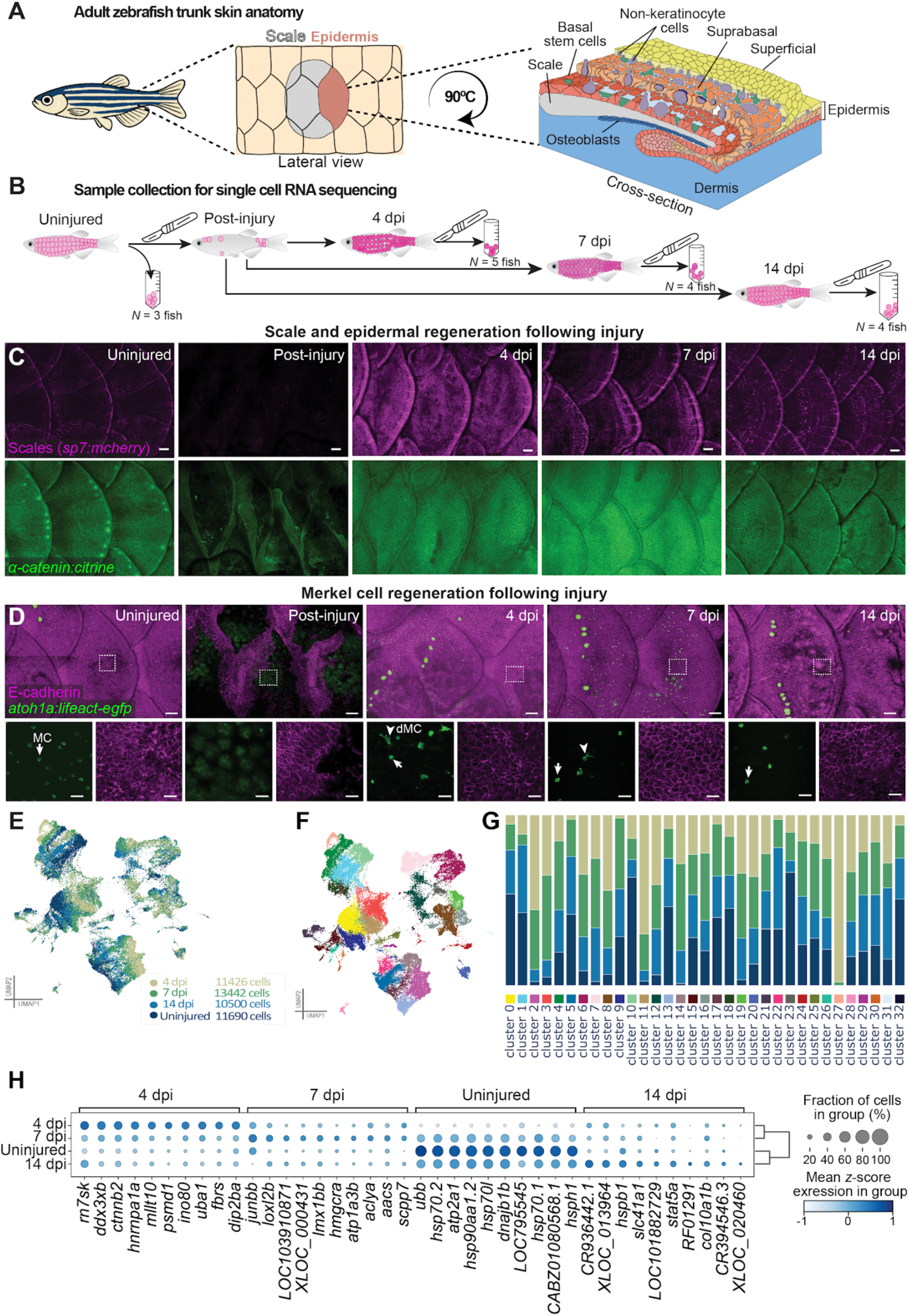
Single-cell RNA sequencing analysis of regenerating skin from adult zebrafish reveals distinct skin cell populations. **A.** Left, Schematic of adult zebrafish trunk skin anatomy. Right, cross section of skin shows dermal scales underneath a stratified epidermis consisting of multiple layers of keratinocytes and non-keratinocyte cells, including MC. **B.** Schematic of homeostatic (uninjured) and regenerating skin sample collection from adult zebrafish trunk at multiple timepoints for single-cell RNA-sequencing. **C.** Representative images of uninjured and regenerating osteoblasts [magenta; *Tg*(s*p7:mcherry*)] and epidermal junctions [green; *Gt(ctnna1-citrine)*] at the indicated timepoints. Scale bars: 100 µm. **D.** Representative images of homeostatic and regenerating epidermis [magenta; *(cdh1-tdTomato)*] and MCs [green; *Tg(atoh1a:lifeact-egfp*)]. Dashed boxes indicated regions shown at higher magnification in lower panels. Post-injury panel shows pigment autofluorescence in the green channel. Arrows indicate MCs; arrowheads indicate dMCs. Scale bars: 100 µm and 20 µm (insets). **E.** Combined UMAP of cells in all the samples based on transcriptomes. Each dot represents a cell colored according to timepoint. Note that each timepoint shows a distinct transcriptional profile. **F.** Unsupervised default clustering of 47,058 skin cells from 4 different samples yields 33 different clusters based on their gene expression profiles. **G.** Sample composition in each cell cluster colored as in (F) shows timepoint-specific enrichment and depletion patterns across cell populations. **H.** Differential gene expression analysis of samples across timepoints. Color represents the mean z-scored expression within each group.

We previously found that scale removal triggers MC regeneration, which involves an initial phase predominated by dMCs that subsequently transition to MCs (Craig et al., 2025). To understand the regenerative dynamics of the MC lineage, we performed scRNA-seq across multiple timepoints following scale removal to enrich for dMCs and MC progenitors. To induce injury and trigger regeneration, we descaled a cohort of fish to remove scales and overlying epidermis (**Fig. 1B**). After descaling, the fish regenerated scales and epidermal cells including MCs as expected (**Fig. 1C,D**). We collected and profiled regenerating skin at 4, 7 and 14 days post injury (dpi) after a second round of descaling (**Fig. 1B**). We selected these timepoints because at 4 dpi the epidermis contains both dMCs and MCs, whereas at 7 and 14 dpi MCs predominate (Craig et al., 2025) (**Fig. 1C,D**). We also collected uninjured skin as a control.

After quality control, we obtained 47,058 cells across four different samples (10,500-13,422 cells/sample). Our scRNA-seq profiling revealed cell populations whose organization varied across timepoints as shown by combined uniform manifold approximation and projection (UMAP) plots (**Figs. 1E, S1A**). Unsupervised Leiden clustering identified transcriptionally distinct subpopulations (**Fig. 1F**), and sample-wise composition analysis of each resulting cluster highlighted dynamic shifts during regeneration (**Fig. 1G**). Gene expression profiles suggested that at the early regeneration timepoints (4 and 7 dpi) the transcriptomic profiles of the cells were distinct from the uninjured sample, but by 14 dpi the transcriptional profile aligned more closely to uninjured cells (**Fig. 1H**). Thus, the global embedding suggests that as skin regenerates the transcriptional profile gradually restores to a homeostatic state over the course of two weeks, similar to previous anatomical observations (Craig et al., 2025; Iwasaki et al., 2018).

To identify the cell types in our dataset, we manually annotated the cell clusters using marker genes described in the literature (**Table S1**) (Aman et al., 2023; Bowman et al., 2023; Brown et al., 2023; Cannon et al., 2013; Carney et al., 2006; Castranova et al., 2025; Dasgupta et al., 2024; Horn et al., 2026; Hou et al., 2020; Kim et al., 2013; Robertson et al., 2023; Zhou et al., 2023). We identified clusters that aligned with expected epidermal and dermal compartments, such as keratinocytes, epidermal non-keratinocytes, and osteoblasts (**Fig. 2A**). Hierarchical clustering illustrated a structured relationship between cell clusters suggesting a higher-order compartment-level organization within the epidermis consistent with our manual annotations (**Fig. 2B**). Differential gene expression (DGE) analysis across the entire dataset captured distinct transcriptional signatures associated with each cell cluster (**Fig. S1B**). These data suggest that, at the molecular-level, trunk skin regeneration is structured along at least three major axes of variation: time, compartment identity and subpopulation heterogeneity.

**Figure 2.**
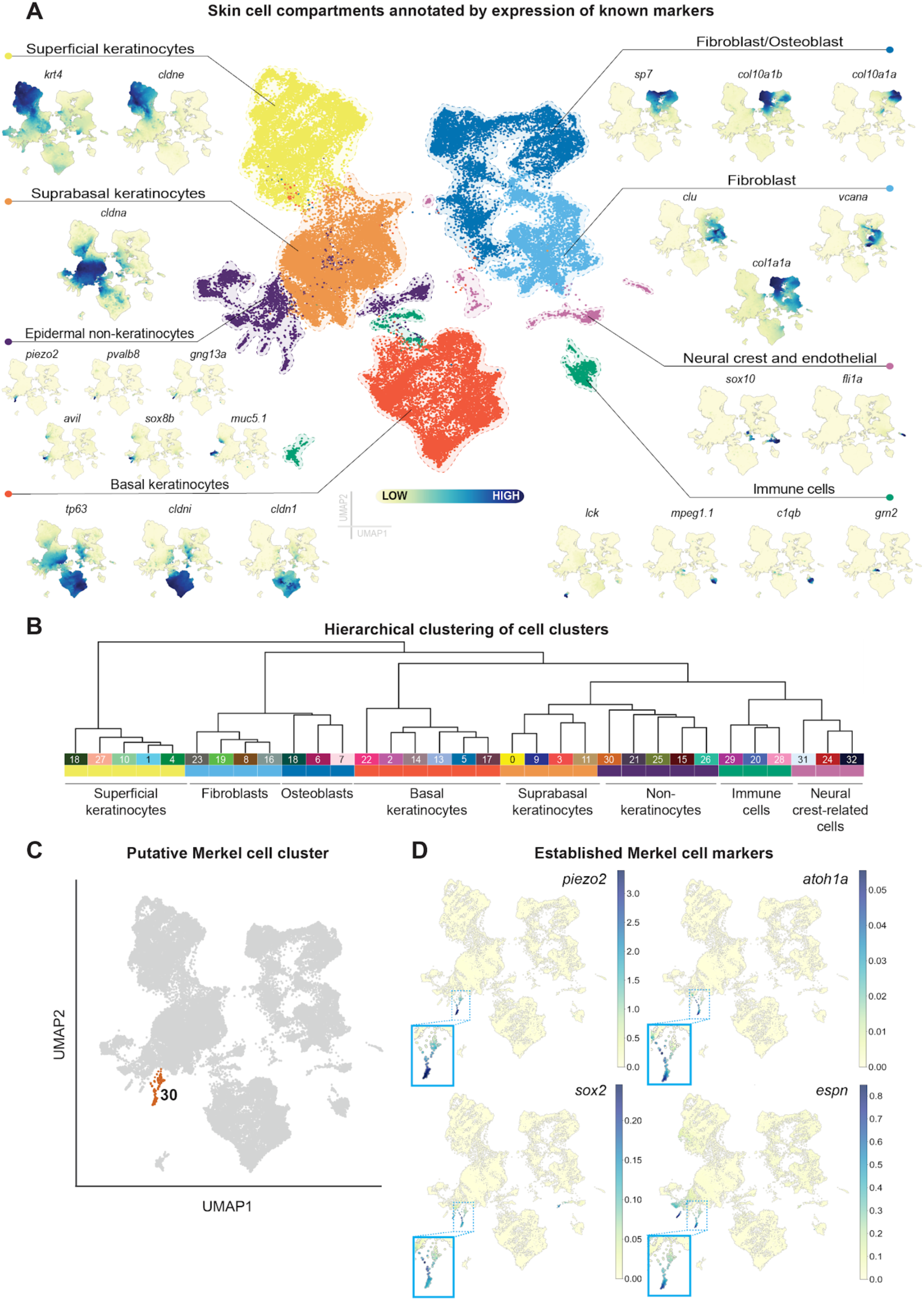
Cluster annotation identifies diverse dermal and epidermal cell types, including Merkel cells. **A.** UMAP plot showing major cell types in skin based on established marker genes (see Table S1 for references). **B.** Hierarchical clustering of cell clusters (colored as in Fig. 1F) labeled according to each major cell type. **C.** UMAP of the entire dataset with cell cluster 30 highlighted. **D.** UMAPs showing imputed expression of established MC markers in the entire dataset. Insets, magnified views of cluster 30.

## Identification of established and novel Merkel cell markers

Among the epidermal non-keratinocytes captured, we found goblet cells, solitary chemosensory cells (types I and II), immune cells and MCs (**Fig. 2A; Table S1**). We identified cluster 30 (**Fig. 2C**; *n*=171 cells) as MCs based on expression of the established MC markers *piezo2, atoh1a, sox2* and *espn (espin)* (**Fig. 2D**) (Brown et al., 2023; Haeberle et al., 2004; Sekerková et al., 2004). Concordantly, gene ontology (GO) analysis of the enriched genes (log_2_ fold change ≥ 0.5 and adjusted *P*-value < 0.05) in cluster 30 revealed a functional signature consistent with MC physiology, including the categories of voltage-gated cation channel activity, transmembrane transporter activity, exocytosis and syntaxin-1 binding (**Tables S2, S3**).

For in-depth analysis of the MC transcriptome, we focused on DGE between the MC cluster and other all clusters, which uncovered enrichment of several genes not previously associated with MCs (**Fig. S2A; Table S2**). Among these novel MC-enriched genes, we concentrated on four implicated in neuronal synaptic function or morphogenesis that encode diverse types of molecules: a δ1-protocadherin [*pcdh9 (protocadherin 9)*], a hexa-EF-hand Ca^2+^-binding protein [*scgn (secretagogin)*], an RNA-binding factor [*rbfox1 (RNA binding fox-1 homolog 1)*] and a single-pass transmembrane protein [*ajap1 (adherens junction associated protein 1)*] (**Fig. 3A**) (Conboy, 2017; Früh et al., 2024; Miozzo et al., 2024; Qin et al., 2020). To validate the expression of these novel markers, we stained regenerating scales from adults expressing an F-actin reporter in MCs [*Tg(atoh1a:lifeact-egfp)*] (Brown et al., 2023) with hybridization chain reaction (HCR) probesets and observed expression of each gene in ≥79% of MCs (**Fig. 3B**). Our dataset thus allows the identification of novel molecules expressed in zebrafish MCs.

**Figure 3.**
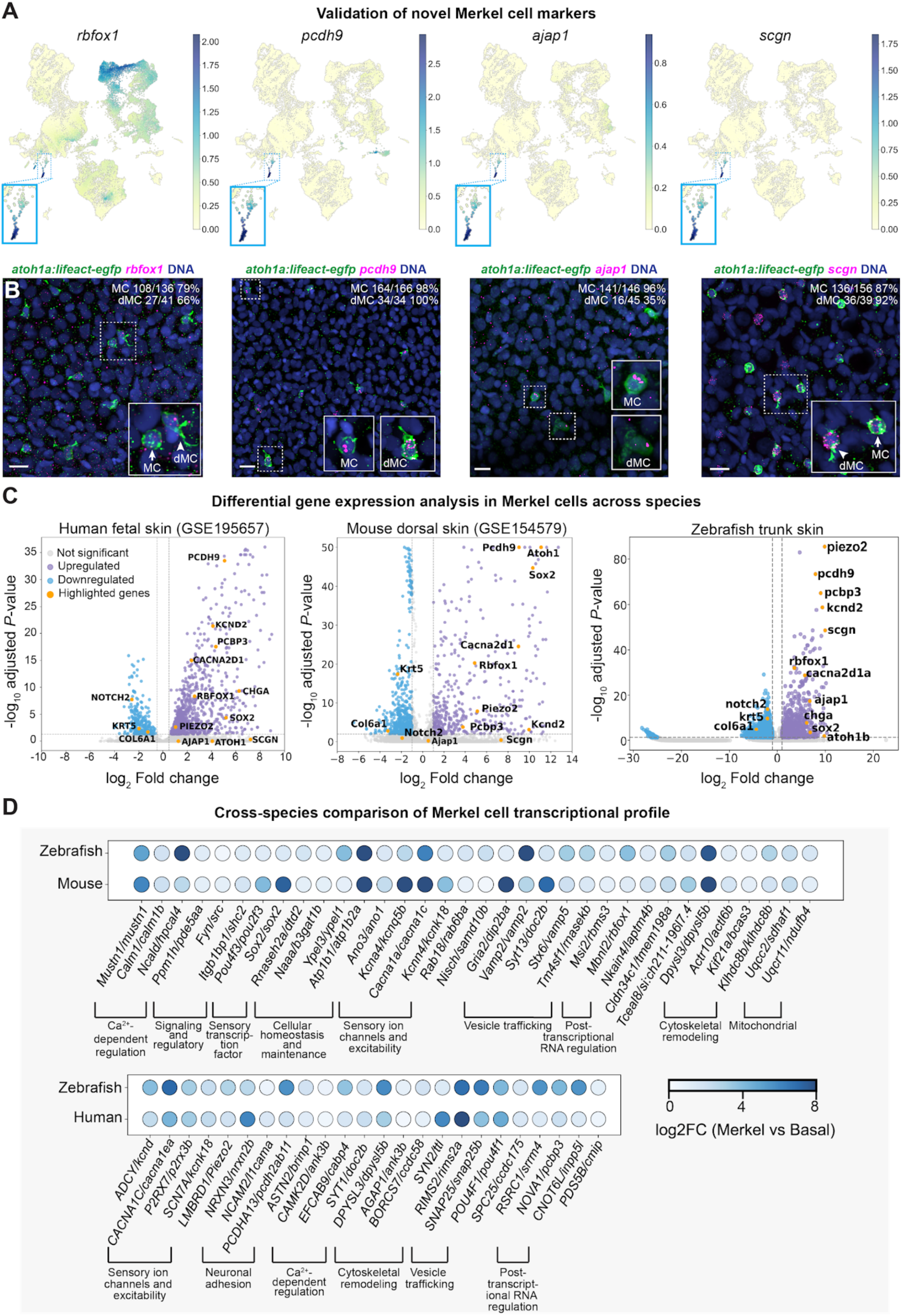
Novel and conserved molecular signatures of vertebrate Merkel cells. **A.** UMAPs showing imputed expression of novel MC markers in the entire dataset. The novel markers were identified by DGE across all epidermal cells (Tables S2, S3). Novel markers are enriched in the MC lineage (cluster 30, Figure 1F). Insets, magnified views of cluster 30. **B.** Representative maximum intensity projections of confocal *z*-stacks of MCs expressing markers identified by scRNA-seq analysis. Each panel shows staining of adult scales expressing a MC reporter [green; *Tg(atoh1a:lifeact-egfp*)] with an anti-GFP antibody and HCR probesets against specific genes (magenta). Arrows indicate MCs, arrowheads indicate dMCs. Labels indicate the numbers of cells positive for each transcript. Cells analyzed from *n* ≥ 8 scales from *N* ≥ 8 adult fish (22-25 mm SL) for each gene in three different experiments. Scale bars: 10 µm. **C.** Volcano plots showing DGEs in MCs relative to non-Merkel epidermal cells from human fetal skin (Glover et al., 2023), mouse dorsal skin (Lin et al., 2020) and adult zebrafish trunk skin (This work). Each point represents an individual gene. Dashed lines denote the log₂ fold-change threshold (0.5), and -log_10_ adjusted *P*-value (0.05) thresholds. Canonical MC markers in all species (e.g., *piezo2*, *sox2*, *kcnd2*) are enriched whereas keratinocyte-associated genes such as *krt5, col5a1*, and *notch2* are depleted. **D.** Dot plots of significantly enriched gene groups in MCs compared to basal keratinocytes between zebrafish and mouse (top) or human (bottom). Dot color represents the average log₂ fold change of MCs relative to basal keratinocytes within each species. Orthologous genes are grouped according to their predicted biological functions. Table S5 lists the genes under each group and their weighted scores.

### Conserved molecular features of vertebrate Merkel cells

To ask to what extent zebrafish MC gene expression is shared with mammalian MCs and to identify a common vertebrate MC transcriptome, we compared our dataset with scRNA-seq of mouse neonatal dorsal hairy skin and single nucleus RNA-seq (snRNA-seq) of human fetal ventral digit skin, both of which captured MCs (Glover et al., 2023; Lin et al., 2020). We compared genes enriched in MCs relative to basal keratinocytes across the three datasets using a threshold of log_2_ fold change ≥ 0.5 and adjusted *P*-value < 0.05. This analysis identified 1,084 enriched genes in human MCs, 627 in mouse and 700 in zebrafish (**Table S4**). MCs in all three species shared enrichment of 35 genes, with an additional 121 shared between human and mouse, 32 between human and zebrafish, and 11 between mouse and zebrafish (**Table S4**). MCs in all three species consistently expressed the mechanotransduction-associated genes *piezo2*, *kcnd2 (potassium voltage-gated channel, Shal-related subfamily, member 2)* and *cacna2d1 (calcium channel, voltage-dependent, alpha 2/delta subunit 1a),* which are required for MC function (Ikeda et al., 2014; Maksimovic et al., 2014; Piskorowski et al., 2008; Woo et al., 2014; Yamada and Gu, 2025). Orthologs of the transcription factor *SOX2* were also enriched across vertebrate MCs, suggesting a conserved MC gene regulatory circuitry (**Fig. 3C, Table S4**). Interestingly, we also identified enrichment of novel MC markers (*rbfox1, pcdh9*) we validated in zebrafish (**Fig. 3C**), suggesting that zebrafish can be used to discover unique features of MCs relevant to mammalian MC biology.

As a complementary approach, we compared the transcriptomes of MCs and basal keratinocytes between zebrafish, mouse and human using SATURN (Rosen et al., 2024). SATURN performs cross-species integration using both protein sequence similarity and gene expression patterns to aggregate functionally similar genes into macrogene modules, thereby facilitating cell type comparison. Principal component analysis (PCA) demonstrated that cell type, rather than species identity, drove transcriptional variance (**Fig. S3A**), further suggesting shared MC transcriptional programs across vertebrates. To quantitatively assess MC transcriptome conservation, we compared SATURN-derived macrogene expression profiles using pairwise Pearson correlation analysis. MCs exhibited moderate to strong positive correlations across species, with the highest similarity between mouse and human (*r* = 0.66), followed by human and zebrafish (*r* = 0.63) and mouse and zebrafish (*r* = 0.31) (**Fig. S3B**). Conserved MC macrogenes included ion channel activity, cell adhesion and cytoskeletal remodeling genes, synaptic membrane components and RNA regulatory factors (**Fig. 3D, Table S5**). These findings indicate deep conservation of both expected and novel MC features that likely play physiologically relevant roles in MCs.

### Transcriptional heterogeneity within the Merkel cell sublineage

To assess transcriptomic dynamics within the regenerating MC lineage, we re-clustered the cells within cluster 30. Unsupervised clustering identified seven MC subclusters (subclusters 0-6) based on transcriptomic profiles (**Fig. 4A**). The sample distribution showed that regenerating MCs predominated in subclusters 2, 4 and 5, whereas subcluster 1 contained the highest fraction of MCs from uninjured skin (**Fig. 4B**).

**Figure 4.**
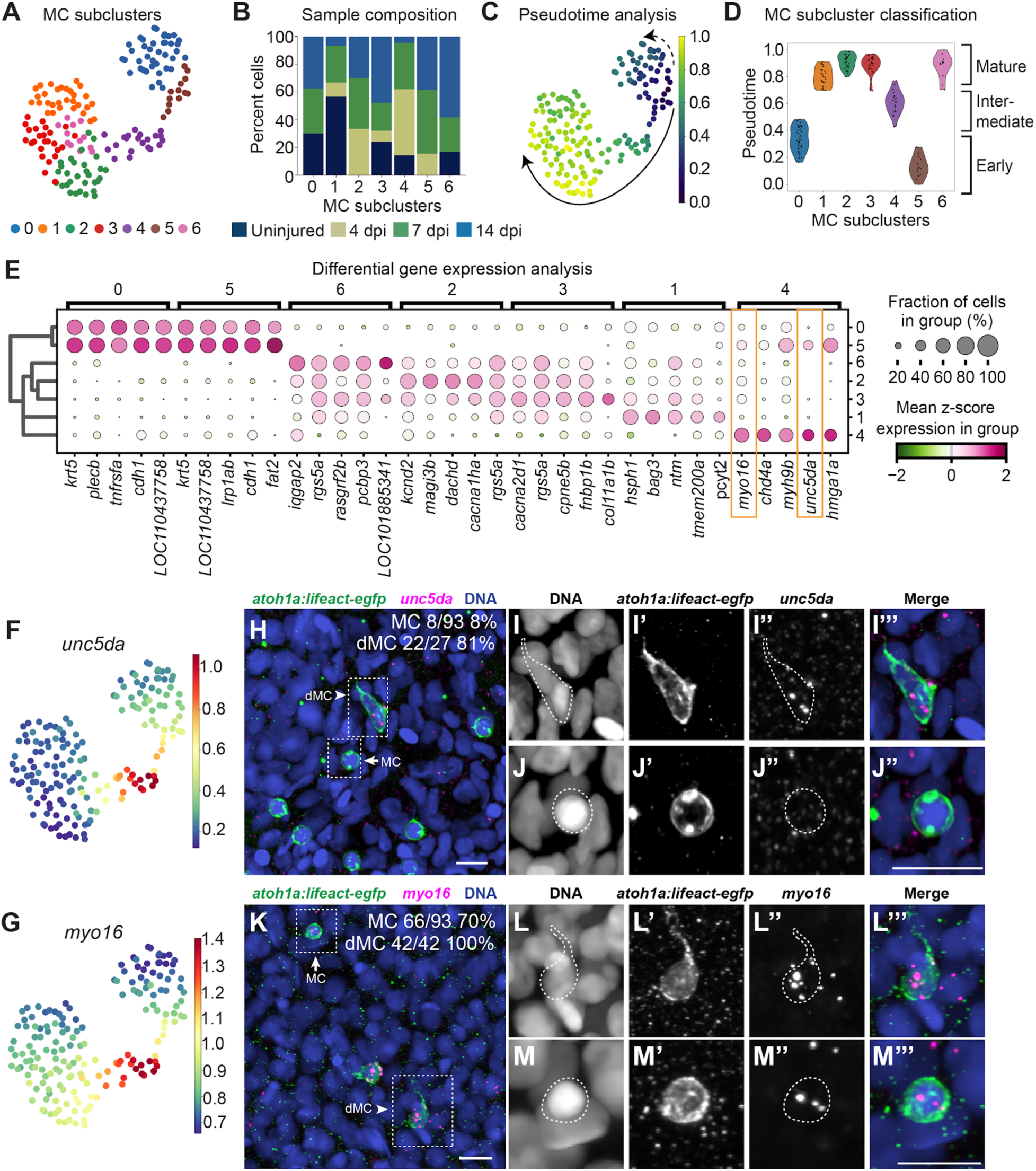
Transcriptional heterogeneity in the Merkel cell lineage. **A.** Unsupervised clustering identified seven subclusters of MCs. Each dot represents a cell. **B.** Sample distribution across the MC subclusters. **C.** UMAP of the MC cluster colored according to a Palantir-generated pseudotime scale of cell state. Dark blue indicates undifferentiated cell states, yellow indicates terminal cell states. Arrows indicate cell trajectories. **D.** Violin plots showing classification of MC subclusters based on their distribution across pseudotime. **E.** Dot plot showing DGE analysis of the top 5 transcripts in each MC subcluster. Size of the dots indicate fraction of cells expressing each gene and the color indicates mean z-scored expression level. For a full list of genes, see Table S6. **F,G**. UMAPs showing imputed expression of the indicated genes in the MC cluster. **H–M’’’.** Representative *z*-projected confocal images of the regenerating scale epidermis at 5 dpi from adult fish expressing a MC reporter [green; *Tg(atoh1a:lifeact-EGFP)*]. Scales were stained with an anti-GFP antibody and HCR probesets against the indicated genes (magenta). Arrows indicate MCs, arrowheads indicate dMCs. *unc5da* staining analyzed from *n*=8 scales from *N*=8 adults (22-25 mm SL) in two different experiments. *myo16* staining analyzed from *n*=5 scales from *N*=5 adults (22-25 mm SL) in two different experiments. Scale bars: 20 µm, insets 10 µm.

To analyze cell state transitions between the subclusters, we used Palantir (Setty et al., 2019) to order cells along a pseudotime axis, which revealed that the subclusters showed a continuous progression of transcriptional states rather than discrete clusters (**Fig. 4C**). Pseudotime analysis identified subcluster 5 as the most undifferentiated state, subcluster 4 as a transitional cell state and subclusters 1, 2, 3 and 6 as the most differentiated (**Fig. 4C,D**). Accordingly, we labeled the subclusters as early progenitors (subclusters 0 and 5), intermediate progenitors (subcluster 4) and mature MCs (subclusters 1, 2, 3 and 6) (**Fig. 4D**).

To identify transcripts characteristic of each MC lineage state, we performed DGE and GO analysis across MC subclusters (**Figs. 4E**, **S4A; Table S6**). We found that early progenitors expressed genes linked to keratinocyte structural integrity, like *krt5 (keratin 5)* and *cdh1 (cadherin 1, type 1, E-cadherin)* (**Figs. 4E, S4A**). By contrast, intermediate progenitors had reduced levels of these epithelial markers with concomitant expression of *sox2* and *atoh1a* (**Fig. S4C,D**) and chromatin binding and regulating factors (**Fig. S4A**), which included *hmga1a (high mobility group AT-hook 1a)* (**Fig. 4E**). As expected, mature MCs showed an enrichment of molecules responsible for ion transport and voltage-gated calcium channel activity, including *kcnd2* and *cacna2d1* (**Figs. 4E, S4A**). Finally, subcluster 1 had reduced expression of ion channel transcripts and instead showed enrichment of protein folding factors, including *bag3 (BCL2 associated athanogene 3)* and *hsph1 (heat shock 105/110 protein 1)* (Chen et al., 2013; Mattoo et al., 2013) (**Figs. 4G, S4A**). As MCs are long-lived cells (Wright et al., 2017), these protein quality control regulators may promote cellular homeostasis.

Based on our *in silico* analysis that subcluster 4 reflected a transitional cell state between epithelial progenitors and MCs, we hypothesized that this subcluster represented dMCs. To test this notion, we generated HCR probesets for genes enriched in subcluster 4. Specifically, we selected *unc5da*, which encodes a transmembrane guidance receptor (Akkermans et al., 2022), and *myo16*, which encodes an unconventional myosin (Bugyi and Kengyel, 2020) (**Fig. 4F,G**). Consistent with our hypothesis, staining for *unc5da* and *myo16* revealed that most or all dMCs expressed *unc5da* (80%) and *myo16* (100%) (**Fig. 4H,I,K,L**). Nevertheless, *unc5da* and *myo16* staining also labeled MCs, albeit less frequently (8% and 70%, respectively) (**Fig. 4H,J,K,M**). Thus, dMCs represent a transcriptionally distinct state within the MC differentiation trajectory associated with expression of *unc5da* and *myo16*.

Finally, as previous work linked *hmga1a* to proliferation of regenerating cardiomyocytes (Bouwman et al., 2025), we performed cell cycle scoring and found that early and intermediate progenitors had positive S or G2M phase scores (**Fig. S4B**), aligning with previous observations that dMCs can divide (Craig et al., 2025). By contrast, most MCs showed negative S or G2M phase scores (**Fig. S4B**), consistent with the post-mitotic nature of mammalian MCs (Merot and Saurat, 1988; Moll et al., 1996; Vaigot et al., 1987; Weber et al., 2023; Wright et al., 2017).

Together, these analyses demonstrate that MC regeneration proceeds through a series of temporally coordinated transcriptional programs. Early steps in the MC lineage involve repression of keratinocyte identity and structural genes, followed by activation of cytoskeletal and chromatin remodeling genes in dMCs and culminating in induction of ion channels associated with MC terminal differentiation.

### Intermediate MC subpopulation has an epithelial-to-mesenchymal transition signature

In contrast with immotile basal keratinocytes and MCs, dMCs display mesenchymal-like behaviors, including motility and dynamic protrusions (Craig et al., 2025). This previous observation led us to ask whether MC subclusters exhibited molecular characteristics of EMT. During EMT, cells express EMT-transcription factors (EMT-TFs), components of EMT-inducing signaling pathways (e.g., transforming growth factor-β/TGF-β) and switch expression of cell adhesion components from epithelial (e.g., E-cadherin/Cdh1) to mesenchymal (e.g., N-cadherin/Cdh2, β-catenin) to gain motility and facilitate invasion (Kim et al., 2002; Kuphal and Bosserhoff, 2006; Sánchez-Tilló et al., 2011). Based on a composite analysis of 24 core EMT genes, which we curated from previous scRNA-seq studies in zebrafish and a literature review (Ansieau et al., 2008; Cano et al., 2000; Comijn et al., 2001; Gao et al., 2015; He et al., 2017; Hou et al., 2020; Kim et al., 2002; Loh et al., 2019; Tang et al., 2022; Tiwari et al., 2013; Verschueren et al., 1999; Xu et al., 2009; Zhang et al., 2016), we identified an EMT transcriptional signature in MC subclusters 4 and 5 (**Fig. 5A**). These clusters showed enrichment for genes encoding EMT-TFs, including *zeb1a (zinc finger E-box binding homeobox 1a)*, *zeb2a* and *zeb2b*; TGF-β pathway components; and N-cadherin and β-catenin (**Fig. 5B**). Notably, cells exhibiting EMT-associated gene expression retained expression of epithelial genes, including *cdh1*, *oclna/b (occludin)*, *tjp1a (tight junction protein 1a)*, *esrp1/2 (epithelial splicing regulatory protein)* (Ikenouchi et al., 2003; Li et al., 2020; Warzecha et al., 2010). This suggests that EMT-related programs initiate without complete loss of epithelial identity. By contrast, epithelial gene expression was reduced in subcluster 4 and remained relatively low across the putative mature MC subclusters (6, 1, 2 and 3), suggesting a sustained reduction in epithelial identity during MC maturation.

**Figure 5.**
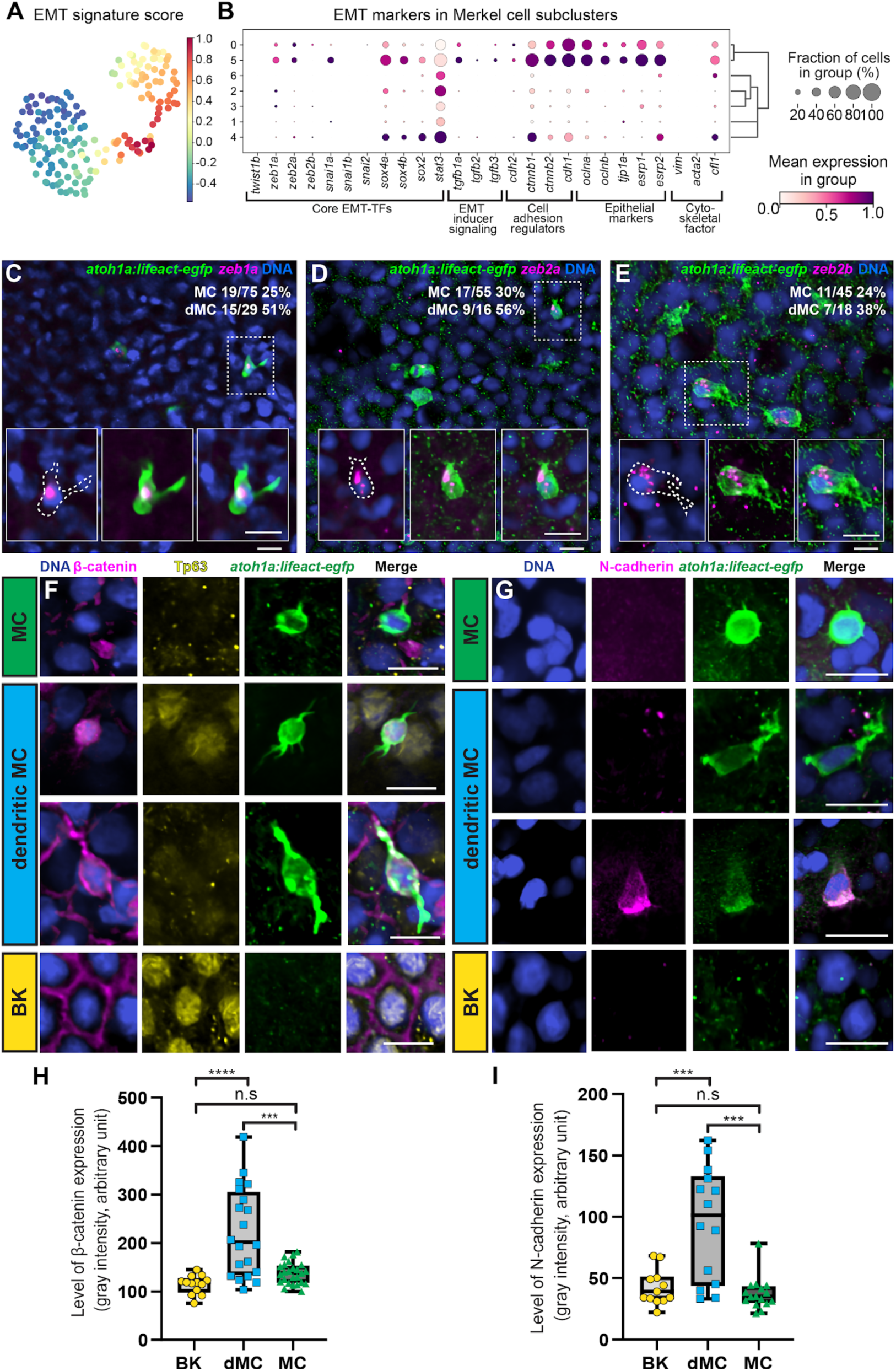
The Merkel cell lineage exhibits an EMT-like signature. **A.** UMAP of the MC cluster colored by the calculated zebrafish EMT score. Higher scores (red) indicate enrichment of EMT-associated gene expression. **B.** Dot plot showing the expression of selected EMT-associated genes across MC subclusters. Dot size represents the percentage of cells expressing each gene within a subcluster, and dot color indicates the mean normalized expression level. Hierarchical clustering identifies subclusters with related EMT-associated transcriptional signatures. **C–E.** Representative single *z*-slice confocal images of the regenerating scale epidermis at 5 dpi from adult fish expressing a MC reporter [green; *Tg(atoh1a:lifeact-EGFP)*]. Scales were stained with an anti-GFP antibody and HCR probesets against the indicated genes (magenta). Cells analyzed from *n*≥5 scales from *N*=5 adult fish (22-25 mm SL) in two different experiments. Scale bars: 20 µm, inset 10 µm. **F,G.** Representative single *z*-slice confocal images of the regenerating scale epidermis at 5 dpi from adult fish expressing a MC reporter [green; *Tg(atoh1a:lifeact-EGFP*)]. Scales were immunostained with an anti-GFP antibody and the indicated antibodies. DAPI (blue) stain indicates DNA. F: 20% of MCs were β-catenin+ in the membrane (5/25), 65% of dMCs with β-catenin+ in the cytoplasm (13/20) and 100% of basal keratinocytes were β-catenin+ in the membrane (13/13). Cells analyzed from *n*=8 scales from *N*=4 adults in two different experiments. G: 0% of MCs were N-cadherin+ (0/16), 43% of dMCs were N-cadherin+ (6/14). No basal keratinocytes expressed N-cadherin (0/13). Cells analyzed from *n*=5 scales from *N*=5 adults (22-25 mm SL) in two different experiments. Scale bars: 10 µm. **H,I.** Quantification of β-catenin and N-cadherin immunofluorescence intensity in basal keratinocytes, dMCs and MCs. Fluorescence intensity (arbitrary units) was measured for individual cells. Each point represents one cell (for H, *n=*13 BK, 20 dMCs and 25 MCs from five fish; for I, *n=*12 BK, 14 dMCs and 16 MCs from seven fish); boxes show the median and interquartile range, and whiskers indicate the minimum and maximum values. A one-way ANOVA with post-hoc Tukey HSD test was used to compare between cell types. \*\*\*\**P* < 0.0001, \*\*\**P* < 0.001; n.s., not significant. BK, basal keratinocytes.

We confirmed the expression of *zeb1a*, *zeb2a* and *zeb2b* in a subset of dMCs and MCs by HCR (**Fig. 5C-E**). Consistent with an EMT-like state, immunostaining for N-cadherin and β-catenin exhibited increased intensity and altered subcellular distribution in dMCs compared to MCs and basal keratinocytes (**Fig. 5F-I**). We conclude that MC progenitors adopt a molecular profile consistent with a partial EMT as they differentiate from a keratinocyte fate.

### Trajectory analysis identifies a candidate population of suprabasal progenitors

To identify upstream progenitors of the MC lineage during regeneration, we tracked cell trajectories *in silico* using Palantir and CellRank (Lange et al., 2022; Setty et al., 2019). To reduce cellular complexity and facilitate trajectory inference, we excluded fibroblasts and osteoblasts followed by unbiased Leiden re-clustering, then performed pseudotime analysis on the remaining epidermal dataset (**Fig. 6A,B**). We re-clustered the remaining cell populations and identified the MC cluster by projecting the original leiden clustering in the newly clustered dataset (**Fig. S5A,B**). Pseudotime calculation identified a basal keratinocyte cluster as the earliest cell state in our epidermal dataset (**Fig. 6B**) and revealed that MCs have a terminal, transcriptionally distinct cluster positioned adjacent to suprabasal keratinocytes within the inferred regeneration landscape (**Fig. 6B**).

**Figure 6.**
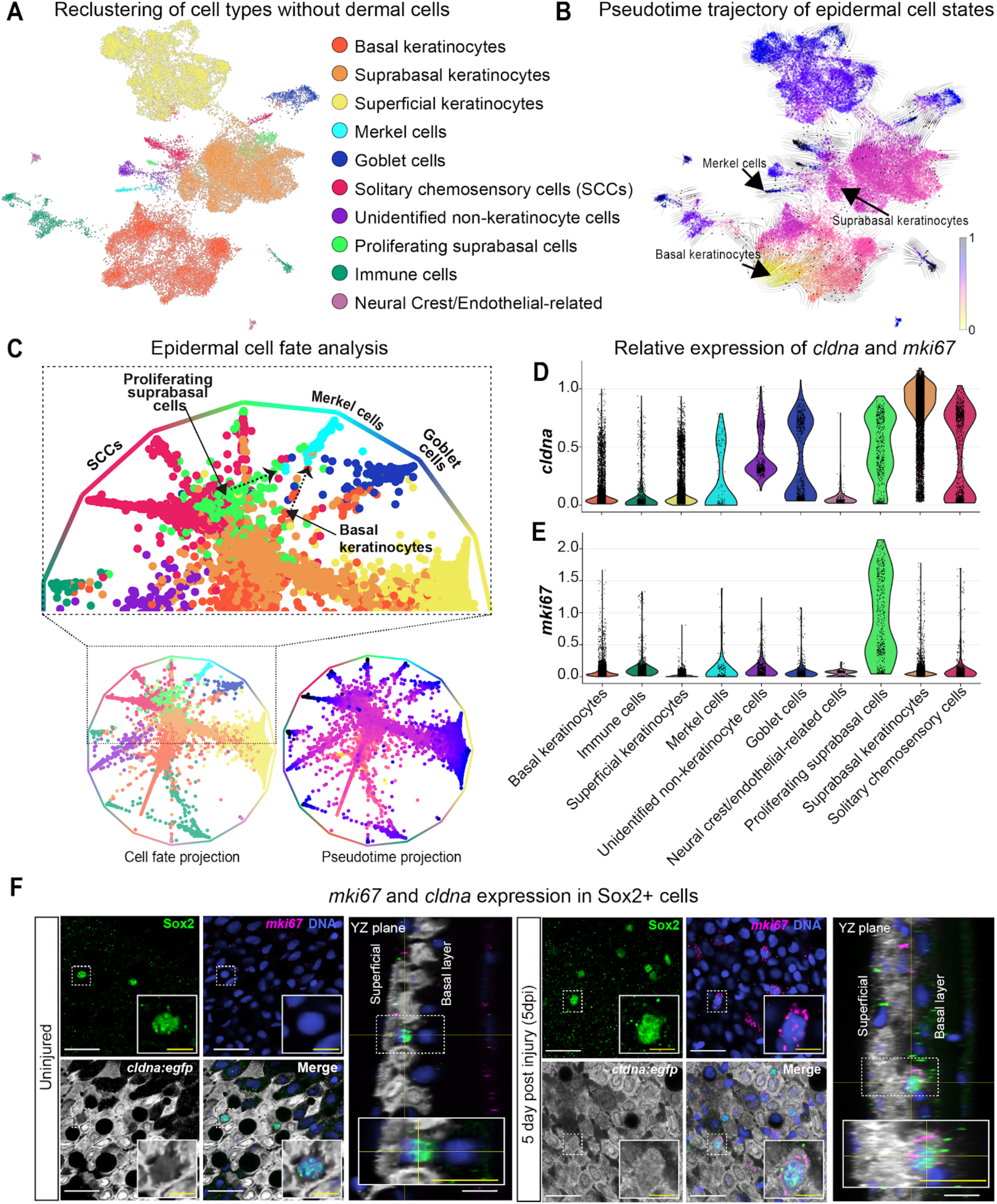
Merkel cell lineage trajectory analysis during regeneration. **A.** UMAP of major epidermal cell types identified by scRNA-seq after exclusion of dermal cells. **B.** Streamline visualization of pseudotime overlaid on the UMAP from A indicating differentiation trajectories. Cells are colored according to pseudotime, with early states shown in yellow and terminal states in blue. Note that basal keratinocytes occupy the earliest pseudotime states, followed by suprabasal keratinocytes, whereas MCs are positioned at the terminal end of the differentiation trajectory. Black streamlines represent the inferred directionality of cellular transitions. **C.** Force-directed graph visualization of epidermal cell states. Annotations illustrate the relative positions of the indicated cell types. Enlarged view (top) highlights the region surrounding the MC lineage. Cell fate projection (bottom left; colored according to cell types in A) identifies putative lineage relationships among epidermal populations, whereas pseudotime projection (bottom right; colored according to pseudotime scale in B) depicts the progression from early to late cell states. Early states shown in yellow and terminal states in blue. Note that the analysis suggests two putative routes (dashed arrows) toward MC differentiation: a direct transition from basal keratinocytes and an alternative trajectory through proliferating suprabasal keratinocytes. **D,E.** Violin plots showing the expression of the indicated markers across major epidermal cell populations. Note that *mki67* expression is highly enriched in proliferating suprabasal keratinocytes, with minimal expression in differentiated epidermal populations, including MCs. **F.** Representative single *z*-slice confocal images of uninjured (top) and regenerating scale epidermis at 5 dpi (bottom) from adult fish expressing *Tg(cldna:egfp*). Scales were stained with anti-GFP (grey) and anti-Sox2 (green) antibodies and HCR probesets against *mki67* (magenta). DAPI (blue) stain indicates DNA.Inset views highlight individual Sox2+ cells (dashed boxes). Orthogonal *yz* sections demonstrate the position of Sox2+ cells. Following injury, Sox2+ cells expressing *Tg(cldna:egfp)* and *mki67* were observed. Arrowhead denotes an elongated nucleus characteristic of dMCs. Scale bars: 20 μm, inset 10 μm.

To further resolve lineage relationships, we analyzed the cell fate trajectories and pseudotime scale of the major epidermal cell types (**Fig. 6C**). Cell fate probability mapping and pseudotime analysis positioned basal keratinocytes at the origin of multiple differentiation trajectories, consistent with their role as epidermal progenitors (Lee et al., 2014) (**Fig. 6C**). One trajectory extended directly from basal keratinocytes toward the MC lineage (**Fig. 6C, right dashed black arrow**). Interestingly, differentiation trajectories also originated from a suprabasal cell subpopulation enriched in *cldna* (*claudin a*), which is broadly expressed in suprabasal keratinocytes (**Fig. 6D, S5C**) (Hou et al., 2020), and *mki67* (*marker of proliferation Ki-67*) (**Fig. 6E, S5D**), a cell proliferation marker (Gerdes et al., 1984), toward several specialized, non-keratinocyte cell types, including MCs (**Fig. 6C, left dashed black arrow**). These observations suggest the existence of two putative routes contributing to MC regeneration.

To investigate a potential relationship between the MC lineage and *cldna+* cells, we began by generating an EGFP reporter line expressed from the endogenous *cldna* locus. *Tg(cldna:egfp)* larvae expressed EGFP in the otic vesicle (**Fig. S6A**), consistent with *in situ* hybridization data (Kollmar et al., 2001). Whole-mount imaging of adults revealed broad but heterogeneous *Tg(cldna:egfp)* expression throughout the trunk epidermis (**Fig. S6B**). Examination of individual epidermal strata showed that ∼80% of cells within suprabasal layers expressed the reporter, whereas a smaller proportion of superficial cells and relatively few basal cells expressed EGFP (**Fig. S6C-E**). To examine transgene expression during skin regeneration, we removed scales and observed strong EGFP signal in suprabasal cells along with an increased percentage of EGFP+ superficial cells at 5 dpi (**Fig. S6D,E**). Thus, *Tg(cldna:egfp)* recapitulates the expected expression pattern and is a novel tool to follow suprabasal keratinocyte dynamics.

We next stained uninjured or regenerating *Tg(cldna:egfp)* skin for *mki67* and Sox2 to label proliferating cells and MCs, respectively. Although we did not detect *mki67* staining in uninjured epidermis (**Figs. 6F, 7A**), cells within regenerating basal and suprabasal layers showed elevated *mki67* expression (**Figs. 6F, 7B**). In uninjured skin, Sox2+ cells had no detectable expression of *Tg(cldna:egfp)* or *mki67* (**Fig. 6F**). By contrast, at 5 dpi, we observed some Sox2+ cells expressing both *Tg(cldna:egfp)* and *mki67* (**Figs. 6F, 7C**) along with cells that expressed only one or neither marker (**Fig. 7D-F**). Previous work demonstrated that nuclear circularity and *z*-depth within the epidermis could differentiate dMCs from MCs, with dMCs having non-circular nuclei and localizing deeper from the surface (Craig et al., 2025). Thus, we quantified nuclear circularity along with the levels of *Tg(cldna:egfp)* and Sox2 expression in Sox2+ cells. Our analysis found that Sox2 expression levels increased substantially in regenerating skin compared to uninjured epidermis (**Fig. 7G**). Additionally, all Sox2+ cells in uninjured skin had circular nuclei (**Fig. 7G’,H**), whereas ∼40% of regenerating Sox2+ cells had non-circular nuclei (**Fig. 7G’,H**). We categorized the putative dMCs characterized by non-circular Sox2+ nuclei into four different subpopulations based on their expression of *Tg(cldna:egfp)* and *mki67* and found that the predominant subpopulation expressed *Tg(cldna:egfp)* but not *mki67* (**Fig. 7I**). In uninjured skin, Sox2+ cells localized just below the superficial layer, whereas all four regenerating subpopulations occupied lower epidermal strata (**Figs. 6F, 7J**), again consistent with a dMC identity. Together, our trajectory analyses and *in vivo* data support a model in which MC regeneration proceeds either through a direct pathway from basal keratinocytes or an indirect pathway involving proliferative suprabasal intermediates (**Fig. 7K**).

**Figure 7.**
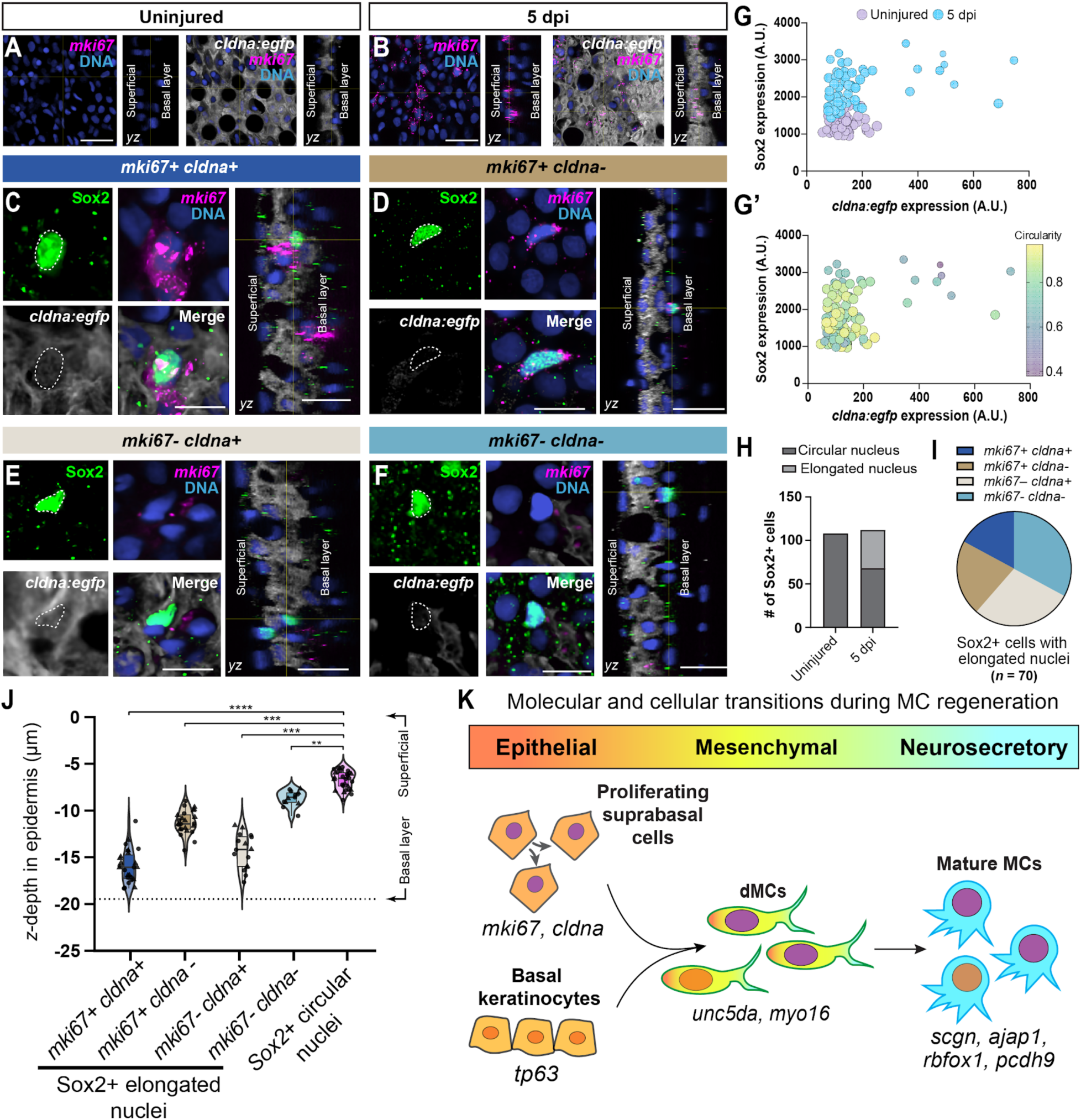
*cldna+* suprabasal cells give rise to regenerating Merkel cells. **A,B.** Representative single *z*-slice confocal images of uninjured and regenerating (5 dpi) adult zebrafish epidermis expressing a *cldna* reporter [*Tg(cldna:egfp)]*. Scales were stained with an anti-GFP antibody (grayscale) and HCR probesets against *mki67* (magenta). Orthogonal views show the localization of proliferating cells within the epidermis. Note that the regenerating epidermis exhibits a marked increase in *mki67*+ suprabasal keratinocytes compared with uninjured tissue. Scale bars: 20 µm. **C–F.** Representative single *z*-slice confocal images of uninjured and regenerating (5 dpi) adult zebrafish epidermis expressing a *cldna* reporter [gray; *Tg(cldna:egfp)*]. Scales were stained with anti-GFP and anti-Sox2 antibodies and HCR probesets against *mki67* (magenta). Orthogonal views show the localization of representative cells within the epidermis. Sox2+ cells were classified according to expression of *mki67* and *Tg(cldna:egfp)* as indicated. Dashed outlines indicate the representative Sox2+ cell shown in each panel. Scale bars: 10 µm, 20 µm (*yz* views). **G,G’.** Scatter plots showing Sox2 and *Tg(cldna:egfp)* expression levels in individual Sox2+ cells from uninjured and 5 dpi epidermis, colored according to sample identity (G) or nuclear circularity (G’). In both plots, point size represents nuclear circularity. Note that Sox2+ cells display a broad range of *Tg(cldna:egfp)* expression levels, indicating heterogeneous characteristics throughout the population. **H.** Quantification of Sox2+ cell nuclear morphology in uninjured and regenerating epidermis. Cells were classified as having either circular or elongated nuclei. **I.** Pie chart showing distribution of the four cellular states among Sox2+ cells with elongated nuclei. **J.** Violin plots of z-depth quantification of Sox2+ cells relative to the superficial epidermal layer. Individual Sox2+ populations occupy distinct positions along the basal-to-superficial axis. The lower dashed line indicates the average depth of the basal surface of basal keratinocytes in the data set (−19.2 µm). Sox2+ cells with elongated nuclei localized at an intermediate position between basal and superficial keratinocytes. Sox2+ cells with circular nuclei occupied more superficial localization compared to cells with elongated nuclei. Violin plots show the distribution of individual measurements; points represent individual cells (*n*=8 scales from 2 fish) and box plots indicate the median and interquartile range. Statistical significance was done using a linear mixed-effects model followed by Tukey-adjusted pairwise comparisons. **, *P* < 0.01, ***, *P* < 0.001, ****, *P* < 0.0001. **K.** Schematic model summarizing the proposed MC lineage during skin regeneration. Lineage analysis suggests that dMCs may arise directly from basal keratinocytes or through a proliferative suprabasal intermediate population (*mki67+cldna*+) (dashed arrows indicate inferred relationships). dMCs exhibit a distinct dendritic morphology and express transitional markers, including *unc5da* and *myo16*, before differentiating into MCs. The gradient indicates a progressive transition from epithelial through mesenchymal-like to neurosecretory cellular states during MC regeneration, with expression of markers characteristic of each state identified in this study listed.

## DISCUSSION

The molecular and cellular trajectory of the MC lineage remains poorly understood partly due to the lack of MC precursors identified in previous scRNA-seq datasets of developing or regenerating mammalian skin. Here, we addressed this knowledge gap by generating a transcriptomic resource of adult zebrafish skin regeneration, encompassing a wide range of dermal and epidermal cell types. We designed our collection scheme to capture regenerating MCs and their progenitors. The higher relative abundance of MCs in our dataset compared to previous sc/snRNA-seq studies in mammalian skin may reflect biological and technical differences—including their more superficial localization within the epidermis, a higher relative cellular density and the relative simplicity of the zebrafish stratified epidermis—which may facilitate more efficient dissociation and capture of MCs in single-cell workflows. Our in-depth analysis of the molecular signature of zebrafish MCs identified novel markers and a core set of genes enriched in MCs across vertebrates. The transcriptomic signature of dMCs revealed an EMT-like signature, characterized by the dynamic expression of cell adhesion molecules and chromatin regulators. Finally, our cell fate trajectory analysis identified a suprabasal keratinocyte→dMC→MC lineage progression during regeneration.

Pioneering studies using either microarrays or bulk RNA-seq cataloged the unique murine MC transcriptome, with an emphasis on the neurosecretory machinery expressed by MCs (Haeberle et al., 2004; Hoffman et al., 2018). Bulk RNA-seq has also served to compare MCs between different murine skin compartments (Nguyen et al., 2019) and analyze POU family gene expression in MCs (Jarvis et al., 2025). A recent scRNA-seq experiment identified an increase in non-coding RNA expression during MC differentiation (Miao et al., 2026). However, a limitation of these previous works is that they relied on fluorescence-activated cell sorting, which biased cell capture and potentially introduced sorter-induced cell stress. By contrast, the dataset generated herein contains a broad representation of dermal and epidermal cell types. Our analysis defined *rbfox1* and *pcdh9* as highly specific markers for MCs within zebrafish skin, which are conserved in humans and mice and expand the possibilities for genetic engineering of MC lineage reagents. Moreover, analysis of the function of these novel genes that represent attributes (e.g., RNA regulation, cell adhesion) poorly understood within the MC lineage will be an exciting area for future study. Our cross-species comparison of the zebrafish MC transcriptome with mammalian datasets revealed a conserved core gene signature, providing strong support for zebrafish as a relevant and tractable model for MC biology. The conserved gene signature included genes encoding the transcription factors Atoh1 and Sox2, and the cation channels Piezo2, Cacna2d1 and Kcnd2, underscoring their roles as fundamental mediators of MC specification and function throughout vertebrate evolution.

To date, the definition of dMCs has relied on cellular morphology with specific molecular markers lacking (Craig et al., 2025; Kim and Holbrook, 1995; Moll et al., 1984; Nakafusa et al., 2006; Narisawa et al., 1993; Tachibana et al., 1997; Tachibana et al., 1998). Through DGE analysis, we identified *myo16* and *unc5da* as genes preferentially enriched in dMCs, which will facilitate the molecular identification and analysis of this transient cell population. Moreover, our scRNA-seq trajectory analysis provides molecular support for the previously proposed model that dMCs serve as direct precursors to MCs (Craig et al., 2025). Our findings suggest that dMCs transiently acquire features of an EMT-like state during MC differentiation. EMT is frequently considered a reversible process, in which cells subsequently undergo mesenchymal-to-epithelial transition (MET) and reacquire epithelial characteristics (Yang et al., 2020). Intriguingly, in our analysis, the resolution of transient mesenchymal features does not appear to involve a transition back to an epithelial state. Epithelial markers that are reduced in dMC-associated subclusters remain transcriptionally low in mature MC subclusters (**Fig. 5B**), whereas genes associated with neurosecretory function become progressively enriched (**Figs. 4E,5B,S4A**). Thus, rather than undergoing a conventional EMT followed by a MET, we speculate that dMCs may use an EMT-like program as a transient intermediate that facilitates departure from their keratinocyte identity and progression toward a neurosecretory fate (**Fig. 7K**). Beyond the skin, the notion that MC precursors have hallmarks of EMT is reminiscent of other examples of sensory cell development from epithelial progenitors that involve differential cell motility and adhesion, such as neuroendocrine cell positioning in the lung (Kuo and Krasnow, 2015) and cell displacements during taste bud assembly (Soulika et al., 2016).

The identification of MC progenitors is of long-standing interest to skin and sensory biologists (Hartschuh et al., 1986). During development, Cre-based lineage tracing indicates that MCs originate from embryonic basal keratinocytes in mice (Morrison et al., 2009; Van Keymeulen et al., 2009) and zebrafish (Brown et al., 2023). Lineage tracing studies during skin regeneration found that touch dome keratinocytes can produce MCs following skin grafts (Woo et al., 2010). Recent work identified Tenascin-C as a marker of touch dome keratinocytes and showed that Tenascin-C+ progenitors can regenerate a fraction of adult MCs (Nguyen et al., 2024). However, human skin contains a paucity of touch domes (Smith, 1970), suggesting that other progenitor populations likely exist. We took advantage of the unbiased collection scheme used to generate our dataset to perform *in silico* lineage tracing, which suggests two potential routes for MC regeneration in zebrafish skin—either directly from basal keratinocytes or via proliferative *cldna+* suprabasal intermediates. Consistent with these *in silico* results, we identified both *cldna+* and *cldna-*Sox2-expressing cells during regeneration. We note that we cannot rule out the decay of fluorophore expression in our experiments using *Tg(cldna:egfp)*. Our observations raise intriguing questions about whether MC development employs both routes and whether each route involves distinct molecular mechanisms. Intriguingly, *cldna+* suprabasal keratinocytes appear to be multipotent progenitors with the potential to generate MCs, other non-keratinocyte cell types and superficial epidermis. Additionally, their proliferative nature suggests they may be self-renewing, however, the detailed *in vivo* analysis of suprabasal keratinocytes and their descendants will require the future development of additional cell type-specific reagents that allow for permanent lineage tracing and inducible cell ablation. Importantly, our work highlights the potential of *cldna*-based transgenics for analysis of suprabasal keratinocytes and their descendants.

There are several limitations to this study. First, our trajectory analysis was restricted to the 171 MC lineage cells captured by scRNA-seq, which, while in line with the relative rarity of MCs within the epidermis, resulted in limited statistical power. Second, our cross-species analysis relied on the limited available mouse and human single cell/nuclear datasets containing MCs. Thus, our analysis likely underestimates the extent of the core vertebrate MC gene expression profile for a variety of reasons including incomplete ortholog annotation and differences in single-cell technologies used, types of skin analyzed and developmental stages. Third, we have not directly investigated the cellular or functional consequences of the observed EMT-associated gene expression. Further mechanistic studies will therefore be required to determine whether these genes actively regulate EMT-like cellular behaviors and define their functional contributions to MC regeneration.

In conclusion, our work characterizes the cellular and molecular progression of regenerating MCs. Human skin injuries often lead to altered sensory perception (Tirado-Esteban et al., 2020) and there is limited evidence for MC regeneration in engineered skin substitutes (Sierra-Sánchez et al., 2021), underscoring that studies of MC regeneration in animal models may lead to improved strategies for regenerative medicine. Moreover, it is tempting to speculate that a deeper understanding of the adult MC lineage may also have implications for deciphering the origins of MCC, given that forced expression of transforming antigens from MC polyomavirus in epidermal lineages can lead to MCC-like phenotypes (Verhaegen et al., 2022; Weber et al., 2026).

## MATERIALS AND METHODS

### Zebrafish

Zebrafish were housed at 26-27°C on a 14/10 h light cycle. Animals of either sex were used in this study. All zebrafish experiments were approved by the Institutional Animal Care and Use Committee at the University of Washington (Protocol #4439-01). Published strains used were: AB (wild type), *(cdh1-tdTomato)^xt18^*(Cronan et al., 2018)*, Tg(atoh1a:lifeact-egfp)^w259Tg^* (Brown et al., 2023), *Gt(ctnna-citrine)^ct3a^* (Žigman et al., 2011) and *Tg(Ola.Sp7:mCherry-Eco.NfsB)^pd46Tg^* [referred to as *Tg(sp7:mCherry)*] (Singh et al., 2012). To control for differences in growth rates, zebrafish post-embryonic development was staged based on SL (standard length). Fish SL was measured using the IC Measure software (The Imaging Source) on images captured on a Stemi 508 stereoscope (Zeiss) equipped with a DFK 33UX264 camera (The Imaging Source).

### Generation of Tg(cldna:egfp)^w2005Tg^

*Tg(cldna:egfp)* was generated by CRISPR-mediated knock-in as previously described (Kimura et al., 2014). A donor plasmid containing the Mbait site, a minimal *hsp70l* promoter, EGFP and *bgh poly(A)* sequences was created using Gibson assembly. The insertion was targeted 170 bp upstream of the endogenous *cldna* start codon using the guide RNA: 5’-TCCACCCTATAAATGTGTCAAGG-3’. The plasmid, Mbait and *cldna* gRNAs, and Cas9 protein were injected into single-cell embryos of the AB strain as previously described (Kimura et al., 2014). Larvae were screened for EGFP expression in otic vesicles at 5 dpf and raised to adulthood and confirmed for skin expression. A founder adult was identified and outcrossed to generate a stable transgenic line designated *Tg(cldna:egfp)^w2005Tg^*.

### Cell dissociation for single-cell RNA-sequencing

16 month old fish of both sexes between 26-31 mm SL were used for trunk epidermal cell dissociation. Individual samples consisted of pooled epidermal cells from 3 uninjured fish, 5 fish at 4 dpi, 4 fish at 7 dpi, and 3 fish at 14 dpi. For regenerating fish, scales were removed from the flanks of fish using an angled dissecting knife (Fine Science Tools 100056-12) and recovered in system water and placed back on the fish facility system until all fish could be processed simultaneously. On the day in which all animals reached the target days post injury (14, 7, or 4 days post injury, and uninjured conditions), scales were removed and placed into separate solutions of 2 ml ice-cold 1x phosphate-buffered saline (PBS)/1% bovine serum albumin (BSA) in a 35 mm dish. PBS was removed and replaced with 2 ml 1x PBS/1%BSA + 1000 u/ml collagenase type I (Gibco 17100-017). Scales were incubated at 37°C for 30 min. Collagenase solution was removed and replaced with 4 ml TrypLE Express (Gibco 12604-013) pre-warmed to 37°C. Scales were triturated 10x with a P1000 every 5 min for a total of 20 min. For the last trituration, a P200 was used. The solution was filtered through a 35 μm filter into a 15 ml conical tube. The dish was washed with 4 ml ice-cold 1xPBS/1%BSA and filtered through the same 35 μm filter into the same 15 ml conical tube. The filtrate was spun at 300 *x* g for 5 min using a swinging bucket centrifuge that was pre-cooled to 4°C. The supernatant was aspirated and resuspended in 1ml of ice-cold 1xPBS/1%BSA. The resuspended pellet was filtered through a 35 μm filter and brought up to a volume of 5ml with 1xPBS/1%BSA. The resuspended solution was moved to a fresh 15 ml conical tube and spun at 300 *x* g for 5 min. After this spin, as much supernatant as possible was removed and the pellet was resuspended in 100 μl of ice-cold 1xPBS/1%BSA. The resuspended pellet was filtered through a 35 μm filter into a 1.5ml centrifuge tube and placed on ice. Live cell viability staining was performed with an aliquot of the cell suspension using calcein-AM and DAPI.

### Single-cell RNA-sequencing library preparation and sequencing

After cell dissociation, cells were fixed, and sequencing libraries were prepared according to the manufacturer’s protocol for whole cells using the Evercode WT v2 kit (Parse Biosciences). Raw FASTQ files from uninjured and 4, 7, and 14 days post-injury (dpi) samples across all Parse sublibraries were demultiplexed using the Parse Biosciences split-pipe (Parse Analysis Pipeline) to generate cell-by-gene count matrices. Reads were mapped to the zebrafish reference genome (GRCz11) and annotated according to the Lawson lab Transcriptome Annotation, V4.3.2 (Lawson et al., 2020), which was modified to replace the *piezo2* (LL0000040575), CU179656.2 (LL0000007848) and LO018508.1 (LL0000013864) gene models with updated *piezo2* isoform annotations from NCBI (XM_021468270.1, XM_021468271.1, XM_017352447.2 and XM_021468277.1). Transcript counts were generated using STAR (Dobin et al., 2013) according to the Parse analysis pipeline.

### Data preprocessing, dimensional reduction and clustering

Single-cell RNA sequencing data were processed and analyzed using Scanpy (v1.11.4) in Python (v3.10.18). Raw count matrices were loaded into AnnData objects and subjected to standard quality control procedures in the scanpy package. Cells with low transcript counts, high mitochondrial gene content or excessive total counts indicative of potential doublets were excluded. A cutoff doublet score of 0.2 was used to eliminate any potential doublets. Cells expressing fewer than ≥ 800 transcripts across the regeneration timepoints and with mitochondrial transcript fractions exceeding 5% were removed. Genes expressed in fewer than 2 cells were also filtered out which allowed us to identify genes with low levels of expression. After quality control filtering, the final dataset contained 31,130 genes for downstream analyses.

Following quality control, counts were then normalized on a per-cell basis using the Scanpy default setting (scanpy.pp.normalize_total) counts and log-transformed using scanpy.pp.log1p. Highly variable genes (HVGs) were identified using scanpy.pp.highly_variable_genes (n_top_genes = 3,500) on a per sample basis. HVGs identified in any of the four ) were combined, resulting in 5,131 unique HVGs, which were used for all downstream analyses to reduce technical noise and improve signal detection.

PCA was performed on the scaled HVG expression matrix to reduce dimensionality. The top 100 principal components were used to construct a k-nearest neighbor (kNN) graph, which was used to compute a UMAP for visualization of the cellular landscape. Clustering was performed using the Leiden algorithm with default resolution parameters to identify transcriptionally distinct cell populations.

### Pseudotime and cell fate mapping

Pseudotemporal trajectory reconstruction and lineage inference were performed using Palantir and CellRank within Scanpy. Following preprocessing and quality control, normalized and log-transformed expression matrices were restricted to HVGs. PCA was performed, and the top 100 principal components were used to compute a kNN graph. Diffusion maps were then calculated to capture the intrinsic manifold structure of the data and model continuous transcriptional transitions. Pseudotime ordering was inferred using Palantir, with basal keratinocytes selected as the root population based on established marker expression and cluster identity. In addition to pseudotime, Palantir was used to estimate cell state entropy, providing a quantitative measure of differentiation potential across the trajectory.

To model lineage commitment and identify terminal cell states, CellRank was applied using a transition probability matrix constructed from transcriptomic similarity and pseudotime-derived directionality.

### Cross-species analysis

Cross-species integration was performed using SATURN, a deep learning–based framework that embeds cells from multiple species into a shared low-dimensional space (Rosen et al., 2024). SATURN leverages protein embeddings derived from the ESM2 language model to infer gene similarity across species and constructs “macrogenes,” defined as groups of functionally related and co-expressed genes conserved across species. Precomputed protein embeddings for human, mouse and zebrafish provided with the package were used in this study.

Prior to integration, annotated cell-by-gene matrices were downsampled to include only MCs and basal keratinocytes to reduce computational complexity. SATURN models were trained using raw (unnormalized) counts with 1,000 macrogenes and 8,000 highly variable genes. Pairwise integrations were performed for each species combination (human–zebrafish, human–mouse, and mouse–zebrafish) using corresponding cell type annotations for MCs and basal keratinocytes, following the recommended workflow. Macrogene embeddings were compared using cosine similarity, and PCA was used to visualize shared and species-specific variation. Gene group expression was filtered by adjusted *P*-value of <0.05.

### EMT-signature score analysis

EMT-associated gene sets were first filtered for genes detected in the MC cluster. Then, for each cell, expression values for curated signature genes were obtained from the MAGIC-imputed expression matrix and standardized across cells using z-score transformation. A signature score was then calculated for each cell as the mean z-scored expression across genes in the curated gene set.

### Scale pluck

A scale pluck protocol was followed according to Craig et al., (2025). Briefly, adult zebrafish were anesthetized in 0.006–0.012% buffered MS-222 (MilliporeSigma, E10521) prepared in system water. Anesthetized fish were positioned on the lid of a Petri dish under a dissection scope. Scales were removed from the trunk using Dumont #5 forceps in a posterior-to-anterior sequence to ensure consistency across samples. Following the procedure, fish were returned to system water and monitored until complete recovery from anesthesia. This protocol was followed to collect samples for HCR and tissue regeneration imaging.

### Adult fish fixation

Adult zebrafish carrying relevant transgenes were euthanized in ice-cold water and fixed in 4% paraformaldehyde (PFA) prepared in 1× PBS at 4°C overnight. Following fixation, samples were washed 3× in 1× PBS to remove residual fixative followed by incubation in DAPI for 10 min. After that, fixed samples were washed 3× in 1× PBS. Then the samples were mounted on 22 mm coverslips within a plastic chamber sealed to the coverslip with vacuum grease. The fish body was secured with molten 1% agarose in system water and the chamber was filled with PBS. The chamber was sealed to a microscope slide with vacuum grease and stored at 4°C until imaging.

### Immunostaining

Zebrafish were anesthetized for 2 min in 0.012% MS-222 prepared in system water. 20-30 scales were plucked from the trunk and transferred in 375 μl of 1× PBS containing 1.5 mL microcentrifuge tubes. Fixation was performed by adding 125 μl of 16% paraformaldehyde (PFA) for a final concentration of 4% PFA incubated for 20 min at room temperature on a gently rotating platform. PFA solution was then removed carefully from the tubes and scales were washed with 500 μl of 0.2% PBS and triton (PBST; 1× PBS supplemented with 0.2% Triton X-100) 3×5 min. Subsequently, blocking solution (10% normal goat serum in PBST, 1× PBS with 0.1% Tween-20) was added to the samples and blocked for 1-2 h at room temperature.

After blocking, samples were incubated with primary antibodies diluted in blocking solution (200 μl total volume). Primary antibodies included mouse monoclonal anti-β-catenin (BD Transduction laboratories, 610153; RRID:AB_397555; 1:500), rabbit anti-Cdh2 (GeneTex, GTX125885; RRID:AB_2885609; 1:500), rabbit polyclonal anti-Tp63 (GeneTex, GTX124660; RRID:AB_11175363; 1:800), chicken anti-GFP (GeneTex, GTX13970; RRID:AB_371416; 1:500) or rabbit anti-Sox2 (GeneTex, GTX124477; RRID AB_11178063; 1:500). Scales were incubated with primary antibodies overnight at 4°C, protected from light. The following day, samples were washed four times for 15 min each in PBST and incubated with secondary antibodies diluted in blocking solution. Secondary antibodies included goat anti-mouse Alexa Fluor 647 (Thermo Fisher Scientific, A32733; RRID: AB_2866492; 1:1000), goat anti-rabbit Alexa Fluor 568 (Thermo Fisher Scientific, A11036; RRID: AB_10563566; 1:1000) or goat anti-chicken Alexa 488 (Thermo Fisher Scientific, A11039; RRID: AB_2534096, 1:1000). Samples were incubated for 2 h at room temperature in the dark, followed by four 15 min washes in PBST. Nuclei were counterstained with DAPI (5 ng/μl; MilliporeSigma, 508741) and washed four times for 5 min each in PBST. Scales were mounted epidermis-side up on glass slides using ProLong Gold (Thermo Fisher Scientific) and cured overnight in dark before imaging.

### Hybridization chain reaction

Custom probesets for *ajap1* (accession XM_021478418.1; set size: 50; amplifier: B3), *pcdh9* (accession XM_009302118.5; set size: 20; amplifier: B3), *piezo2* (accession XM_021468270.1; set size: 20; amplifier: B3), *mki67* (accession NM_001277446.2; set size 104, amplifier: B3), *myo16* (accession XM_073912466.1; set size 50, amplifier B1), *rbfox1* (accession XM_068216714.2, set size: 42, amplifier B3), *scgn* (accession NM_001005776.1; set size: 22; amplifier: B1), *unc5da* (accession NM_001328349.2; set size: 33; amplifier B3), *zeb1a* (accession XM_073929440.1, set size 40, amplifier: B3 ), *zeb2a* (accession NM_001114551.1, set size: 40, amplifier: B3) and *zeb2b* (accession XM_073953097.1, set size: 40; amplifier: B3) were designed using insitu_probe_generator (Kuehn et al., 2022). HCR on adult zebrafish scales was performed as described previously (Brown et al., 2023).

### Confocal imaging

Confocal z-stacks were collected using a A1R MP+ confocal scanhead mounted on an Ni-E upright microscope (Nikon) using a 10× air objective for whole fish imaging and 25× water dipping or 40× oil immersion objective for fixed image acquisition. Images were acquired in galvo scanning or resonant scanning mode and post-processed using the denoise.ai function in NIS-Elements (Nikon).

### Image analysis

Image processing was performed using FIJI/ImageJ (Schindelin et al., 2012). Some images were *z*-stacked and displayed as maximum intensity projections. For protein level quantitation, confocal *z*-stacks were 3D projected and subjected to thresholding in ImageJ using Huang’s algorithm to outline *Tg(atoh1a:lifeact-egfp)+* and Tp63+ cells. Fluorescence intensity was measured as the mean gray value within the outlined cells using the ‘Analyze Particles’ feature of ImageJ. Fluorescence intensity values were normalized first by subtracting mean background fluorescence and then to the minimum and maximum mean gray value among all the cells.

### Statistics

Statistical tests used in these studies are listed in individual figure legends. Plots were created using Graphpad (Prism), R or Python. *N* refers to the number of individual fish, and *n* refers to the number of scales used in different experiments. For protein quantitation, we performed two independent experiments consisting of scales from at least 5 fish including both male and female sex.

## ACKNOWLEDGEMENTS

We thank the LSB Aquatics staff for animal care. We thank the lab of Dave Raible for providing the Mbait donor plasmid. The authors are grateful to all members of the Rasmussen lab for discussion, technical assistance, and support.

## AUTHOR CONTRIBUTIONS

Conceptualization: A.S.F., J.P.R., E.J.A.Q.; Methodology: A.S.F., E.P.; Software: A.S.F, E.J.A.Q.; Validation: A.S.F., S.M.S., E.C.B., S.M.D.; Formal Analysis: A.S.F., E.J.A.Q.; Investigation: A.S.F., E.P., S.M.S., E.C.B., S.M.D.; Resources: E.P.; Data Curation: A.S.F.; Writing – Original Draft: A.S.F., J.P.R.; Writing – Review & Editing: A.S.F., J.P.R.; Visualization: A.S.F., E.J.A.Q; Supervision: A.S.F., J.P.R.; Project Administration: J.P.R.; Funding Acquisition: E.C.B., S.M.S., J.P.R.

## COMPETING INTERESTS

No competing interests declared.

## FUNDING

This work was supported by an Institute for Stem Cell and Regenerative Medicine Graduate Fellowship to E.C.B., the Cell and Molecular Biology Training Grant from NIGMS [T32 GM136534] to S.M.S., and the National Institutes of Health [R01 HD107108] to J.P.R..

## DATA AND RESOURCE AVAILABILITY

### Lead contact

Request for further information and resources should be directed to and fulfilled by the lead contact, Jeffrey P. Rasmussen.

## Material availability

Plasmids and transgenic fish generated in this study are available upon request.

## Data and code availability

Accession numbers will be finalized before journal publication.

## SUPPLEMENTARY INFORMATION

### Supplementary Figures and Figure Legends

**Figure S1.**
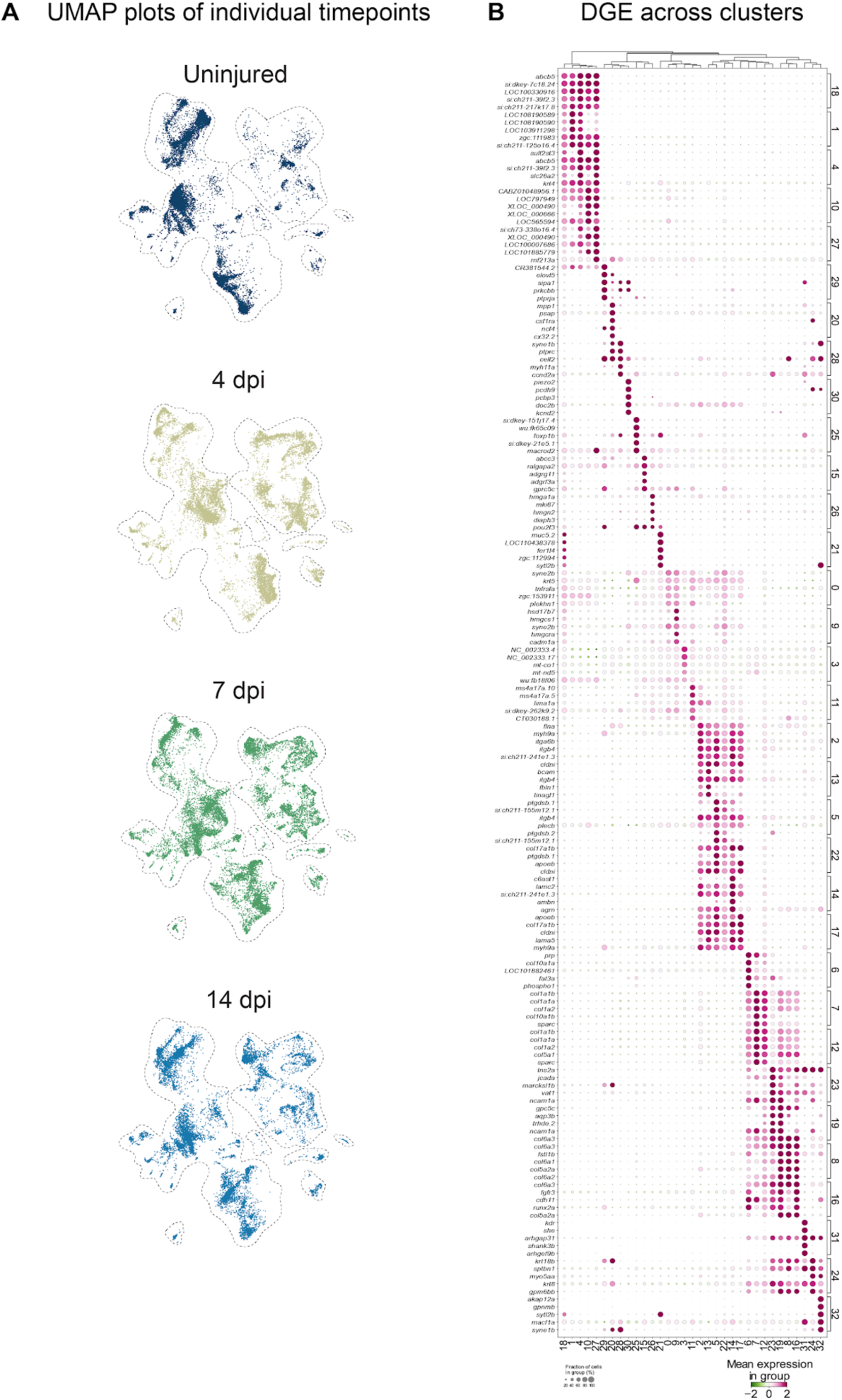
Single-cell transcriptomic landscape of the regenerating zebrafish trunk skin. **A.** UMAPs of all cells collected in each sample. **B.** Dot plot of the top 5 differentially expressed marker genes across clusters identified in Figure 1F. Cell populations (columns) and marker genes (rows) are hierarchically clustered based on gene expression patterns. Dot size represents the percentage of cells within each cluster expressing a given gene, whereas dot color indicates the mean z-scored expression. Distinct transcriptional signatures define each cell population.

**Figure S2.**
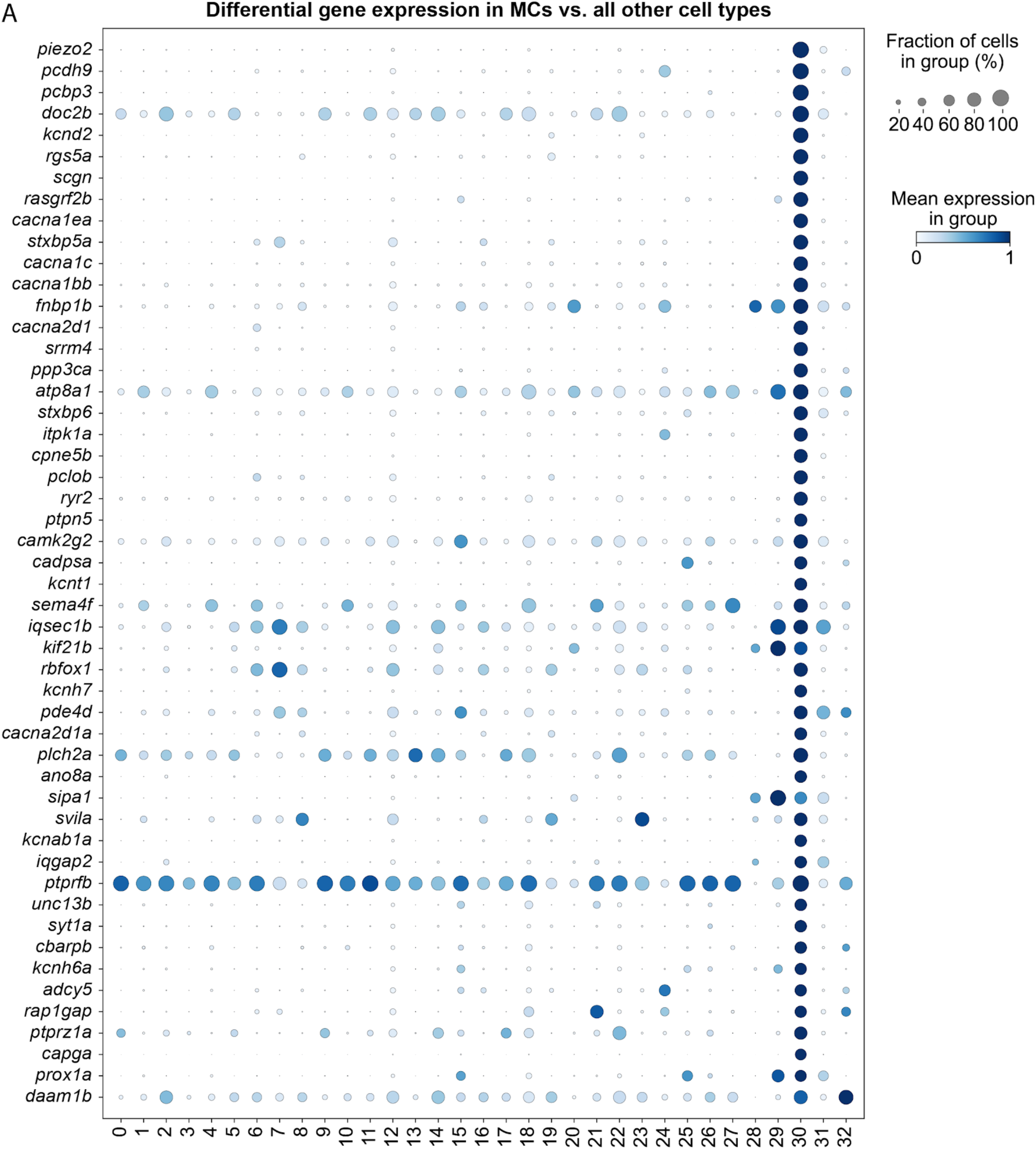
Expression of Merkel cell-enriched genes across skin cell clusters. **A.** Differential gene expression identifies molecular markers of zebrafish MCs. Dot plot showing the expression of top 50 genes significantly enriched in MCs relative to all other skin cell populations identified by scRNA-seq. Dot size represents the fraction of cells expressing each gene, and dot color indicates the scaled mean expression level. The analysis confirms enrichment of established MC markers (*piezo2*, *sox2*) and identifies additional MC-enriched genes. For a complete list, see Table S2.

**Figure S3.**
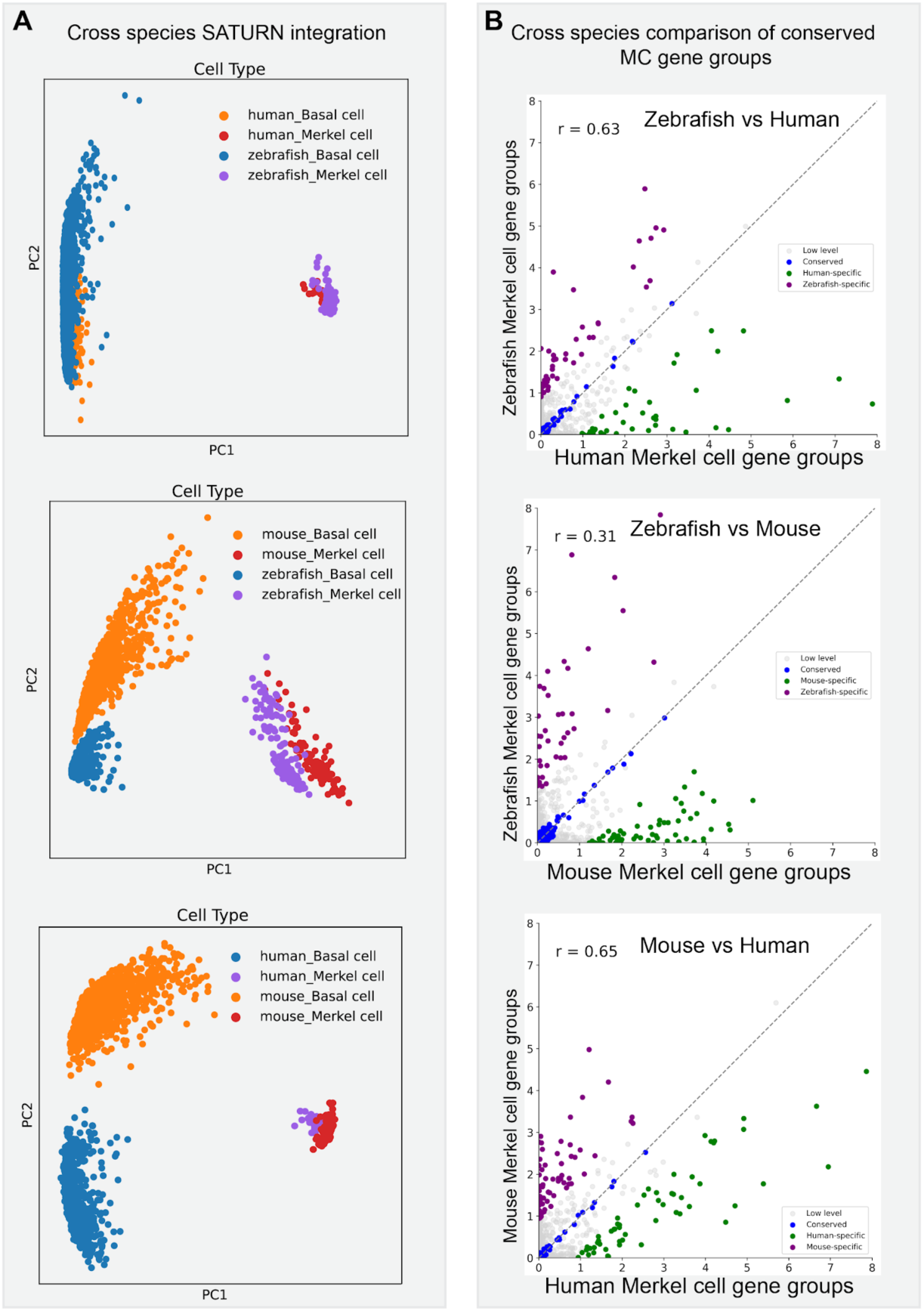
Cross-species transcriptomic integration and conservation analysis of Merkel cells. **A.** Principal component analysis (PCA) of SATURN-generated gene group (macrogene) embeddings comparing zebrafish, mouse and human cell populations. Pairwise integrations of zebrafish-human (top), zebrafish-mouse (middle), and human-mouse (bottom) demonstrate that MCs from different vertebrate species cluster together and separate from basal keratinocytes, indicating conservation of the MC transcriptional program across species. **B.** Cross-species comparison of MC-enriched gene groups (macrogenes) identified by SATURN. Scatter plots compare the expression of macrogene groups between zebrafish and human (top), mouse and human (middle), and zebrafish and mouse (bottom). Each point represents a macrogene group, with colors indicating conserved gene groups (blue), species-specific gene groups (green and purple) or lowly expressed groups (gray). Pearson correlation coefficients (r) are indicated for each comparison, demonstrating the highest transcriptional similarity between mouse and human MCs, while zebrafish MCs retain a substantial conserved molecular signature with both mammalian species.

**Figure S4.**
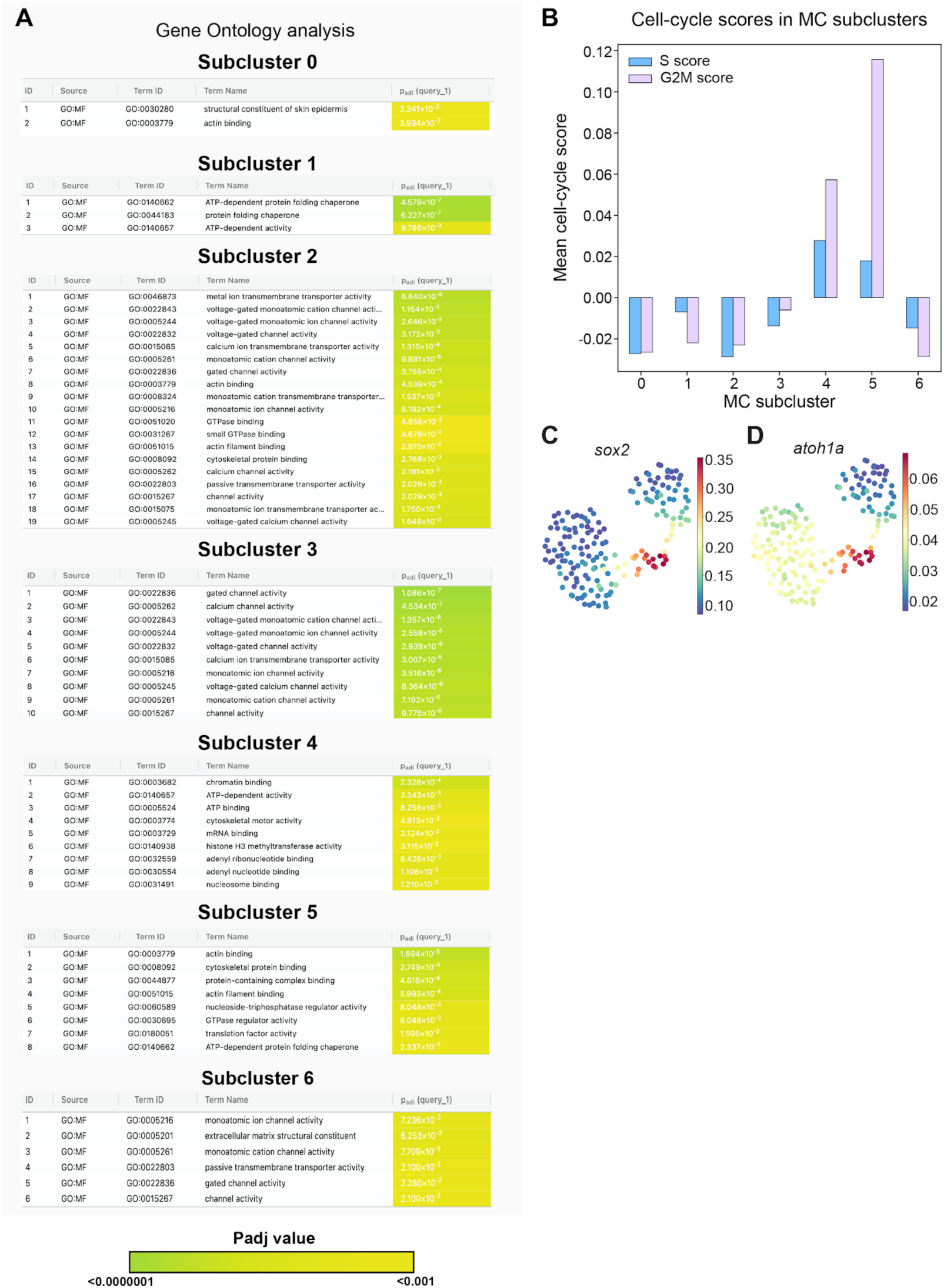
Molecular features of transcriptionally distinct Merkel cell subclusters. **A.** Gene Ontology enrichment analysis using gProfiler (https://biit.cs.ut.ee/gprofiler/gost) for each MC subcluster. Representative significantly enriched biological processes for each cluster are shown. Adjusted *P*-values are indicated by the color scale. **B.** Barplots showing MC subclusters with predicted cell cycle scores (S and G2M score). **C,D.** UMAPs of expression of the indicated genes, color bars indicate level of expression. Each dot represents a cell.

**Figure S5.**
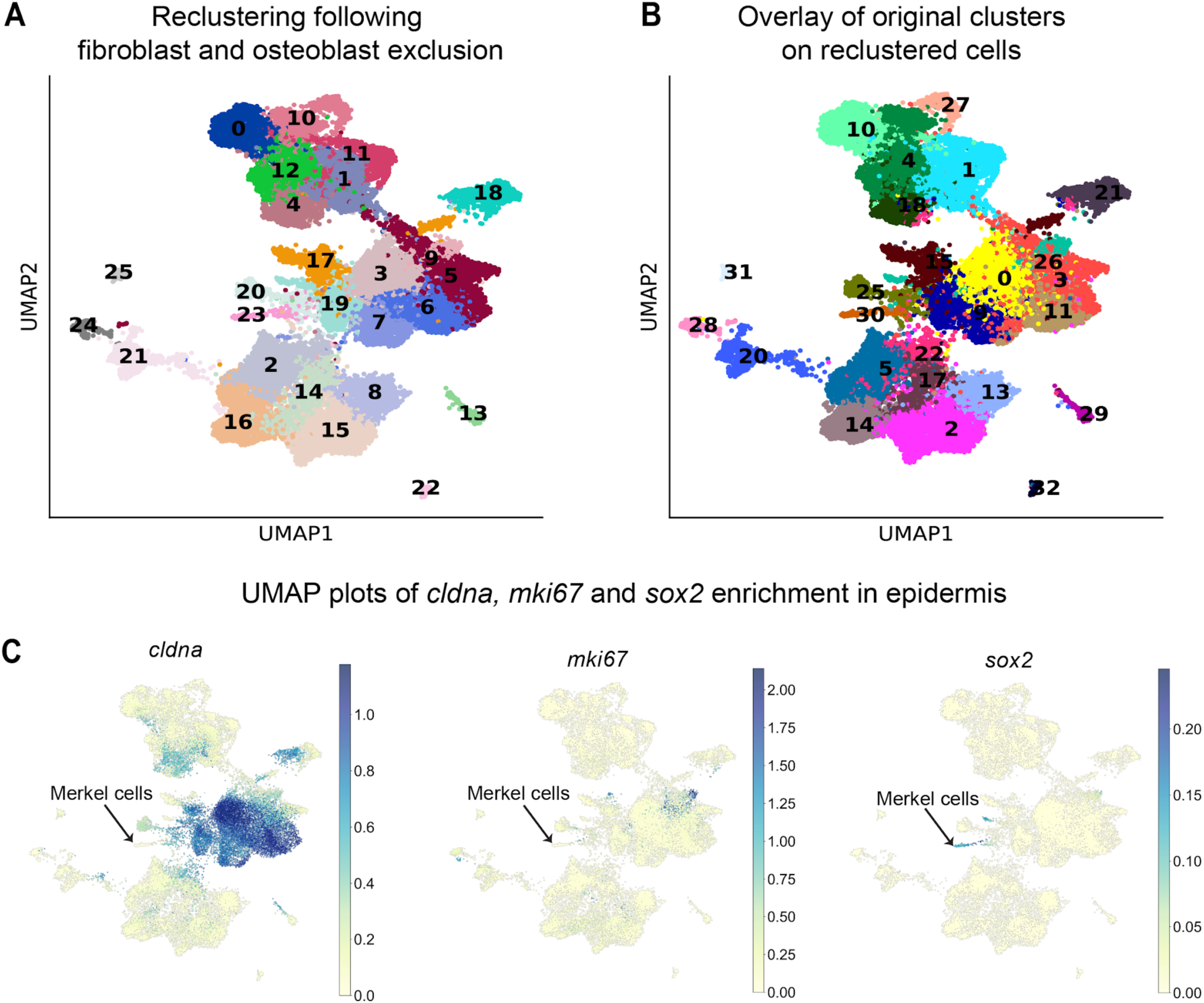
Reclustering of epidermal cells following dermal exclusion. **A.** UMAP of unsupervised Leiden clusters of the reclustered scRNA-seq dataset following exclusion of osteoblasts and fibroblasts. **B.** UMAP of the reclustered scRNA-seq dataset overlaid with the original cell clusters from Figure 1F. **C.** UMAPs of expression of the indicated genes across the reclustered scRNA-seq dataset. Color bars indicate level of expression; each dot represents a cell.

**Figure S6.**
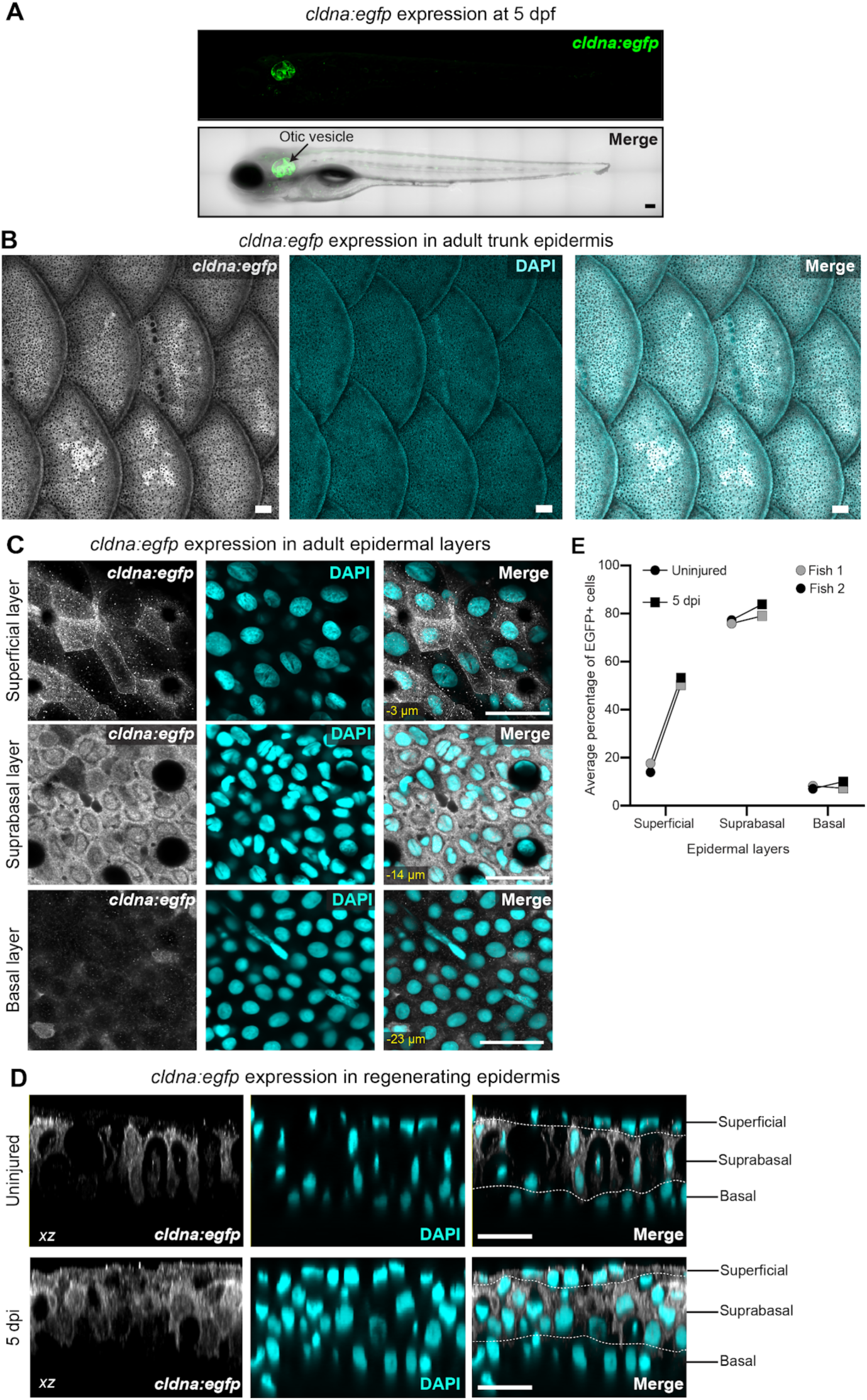
Characterization of the *Tg(cldna:egfp)* expression pattern. **A.** Representative confocal image of *Tg(cldna:egfp)* expression in a 5 day post-fertilization (dpf) larva. Top, maximum intensity projection of *Tg(cldna:egfp)* fluorescence (green). Bottom, EGFP expression overlaid on the transmitted-light channel (merge), showing enriched expression in the otic vesicle. Scale bar, 100 μm. **B.** Representative confocal images of *Tg(cldna:egfp)* expression in the adult trunk epidermis (20.7 mm SL) showing *Tg(cldna:egfp)* expression (gray), DAPI (cyan) and merged channels. Scale bars, 100 μm. **C.** Representative single *z*-slice of superficial, suprabasal and basal epidermal layers showing the distribution of *Tg(cldna:egfp)* expression. DAPI labels nuclei (cyan). Relative z-position of each slice to the superficial layer is indicated in the merge panel in yellow. Scale bars, 20 μm. **D.** Orthogonal views of the adult *Tg(cldna:egfp)* epidermis in the indicated conditions. Images show *Tg(cldna:egfp)* (gray), DAPI (cyan) and merged channels. Superficial and basal boundaries are indicated. Scale bars, 20 μm. **E.** Quantification of the percentage of EGFP+ cells within superficial, suprabasal and basal epidermal layers in uninjured and 5 dpi tissue. Points represent the average percentage of EGFP+ cells calculated from ≥ 8 different scales from 2 fish; the same fish were analyzed under uninjured and regenerating conditions. Lines connect measurements from the same fish.

## Supplementary Tables

**Table S1. Marker genes used for cell type identification. - xlsx file**

**Table S2. Differentially expressed genes in zebrafish Merkel cells compared with other skin cell types. - xlsx file**

**Table S3. Gene ontology enrichment analysis of zebrafish Merkel cells. - xlsx file**

**Table S4. Cross-species comparison of Merkel cell-enriched genes. - xlsx file**

**Table S5. Conserved Merkel cell-associated macrogenes identified in SATURN analysis. - xlsx file**

**Table S6. Differentially expressed genes among zebrafish Merkel cell subclusters. - xlsx file**

## Notes

### Competing Interest Statement

The authors have declared no competing interest.

